# EEG-Based Focus Estimation Using Neurable’s Enten Headphones and Analytics Platform

**DOI:** 10.1101/2021.06.21.448991

**Authors:** Ramses Alcaide, Nishit Agarwal, Jegan Candassamy, Sarah Cavanagh, Michelle Lim, Benyamin Meschede-Krasa, James McIntyre, Maria V. Ruiz-Blondet, Brandon Siebert, David Stanley, Davide Valeriani, Ali Yousefi

## Abstract

We introduce Neurable’s research on focus using our recently developed Enten EEG headphones. First we quantify Enten’s performance on standard EEG protocols, including eyes-closed alpha rhythms, auditory evoked response and the P300 event-related potential paradigm. We show that Enten’s performance is on-par with established industry-standard hardware. We then introduce a series of experimental tasks designed to mimic how focus might be maintained or disrupted in a real-world office setting. We show that (A) these tasks induce behavioral changes that reflect underlying changes in focus levels and (B) our proprietary algorithm detects these changes across a large number of sessions without needing to adjust the model per participant or recording session. Through manipulation of our experimental protocol, we show that our algorithm is not dependent on gross EMG artifacts and it is driven by changes in EEG. Finally, we evaluated the model’s performance on the same subject across several days, and show that performance remained consistent over time. Our model correctly captured 80% ± 4.1% of distractions present in our experiments with statistical significance. This indicates that our model generalizes across subjects, time points, and conditions. Our findings are based on EEG data collected from 132 participants across 337 sessions and 45 different experiments.

## 1 Introduction

Brain-computer interfaces (BCIs) have historically been successful in providing mediums of communication and control to individuals with severe motor impairment^1^. These successes, along with technological advances in BCIs, have expanded the opportunities for an average person to use BCIs in everyday life. In particular, modern life and the diversity of individual needs has pushed BCI research towards enhancing individual productivity, mental health, and relaxation. Still, there remain a number of scientific, technological, and engineering hurdles that need to be solved. Over the last 5 years, Neurable has been building an electroencephalography (EEG) headset (Enten) that can robustly estimate focus. Here, we demonstrate the challenges we faced and advances we made in this domain to build Enten and its corresponding analytics platform.

There are many challenges in building an everyday BCI. For example, most existing BCI applications are created within a controlled laboratory environment and fail to maintain their effectiveness in an unconstrained environment. Most BCI products in the market are either obstructive or come with limited capability, which hinders their actual benefit in an individual’s daily life. At Neurable, we have introduced Enten, an EEG headset which we believe will overcome some of the limitations of previous EEG headsets. Not only is it designed to be comfortable enough for daily use, but it also records high-quality neural activity that can track focus level during the day. Enten is also equipped with many other utilities and sensors, including accelerometer, gyroscope, bluetooth, microphones and speakers. This hardware helps us to build an inclusive picture of human brain dynamics, biological signals, and their interaction with the surrounding environment. We argue Enten is the first generation of EEG-based BCI satisfying most of the characteristics of the ubiquitous BCI.

The challenge in developing a daily use BCI is not limited to device development. Deriving a proper inference from neural data is a big hurdle. Neural data is often buried in noise, movement artifacts, and more importantly, multimodal aspects of brain dynamics making it difficult to isolate the original signal. To address this challenge, we developed several cognitive experiments to build a better picture of human behavioral changes in settings of focus and distraction. Neural and behavioral data captured in these experiments were extensively analyzed and used in the development of our analytics platform. Here, we describe our experimental design and provide information on the extent of our experiments built over eight years of research and conducted in the last two years at Neurable. We discuss how an individual’s behavior shifts in response to a distraction, and how those changes are linked to focus. We explore possible neural correlates that encode focus-related behavioral changes, as well as how these neural features are utilized in our machine learning (ML) analytics platform to estimate focus. The fundamental question is whether our measure of focus is robust to the presence of artifacts imposed by an unconstrained, non-laboratory setting. To address this question, we designed a set of experiments with controlled levels of muscle and movement artifacts to better study resilience and repeatability of our measure of focus. These collective experiments let us study how artifacts impact our neural measure of focus and how those artifacts are removed through our analytical platform.

In this study, we first verify that Enten captures neural activity in a set of standard EEG protocols and compare our findings with industry-standard EEG headsets. Next, we review different cognitive experiments that we have developed to study human behavior during periods of distraction and focus. We demonstrate evidence of consistent changes in behavior in response to those distractions and present possible neural correlates of distraction and focus on individual and population levels. We discuss our neural-based measure of focus and show its performance results in a cohort of experiments and settings. We further expand these analyses in experiments designed to replicate daily-life settings with different noise and artifact elements to show that our analytical platform is robust to the presence or absence of artifacts in its estimation of focus. Finally, we provide evidence for stability of our analytics measure via a longitudinal analysis.

## 2 Overview of Enten EEG hardware

Enten is a Bluetooth around-ear headphone with 20 electrodes placed around the ear pads and records EEG signals at sampling rate of 500 Hz and 24-bit resolution. Our electrodes are made of a conductive mesh that wraps around the ear pads. Each electrode makes use of built-in active amplification. Figure 1 shows the headset and electrode locations. Enten also includes active noise-cancellation, high-quality audio and microphones, and the battery lasts 12+ hours on a single charge.

**Figure 1.**
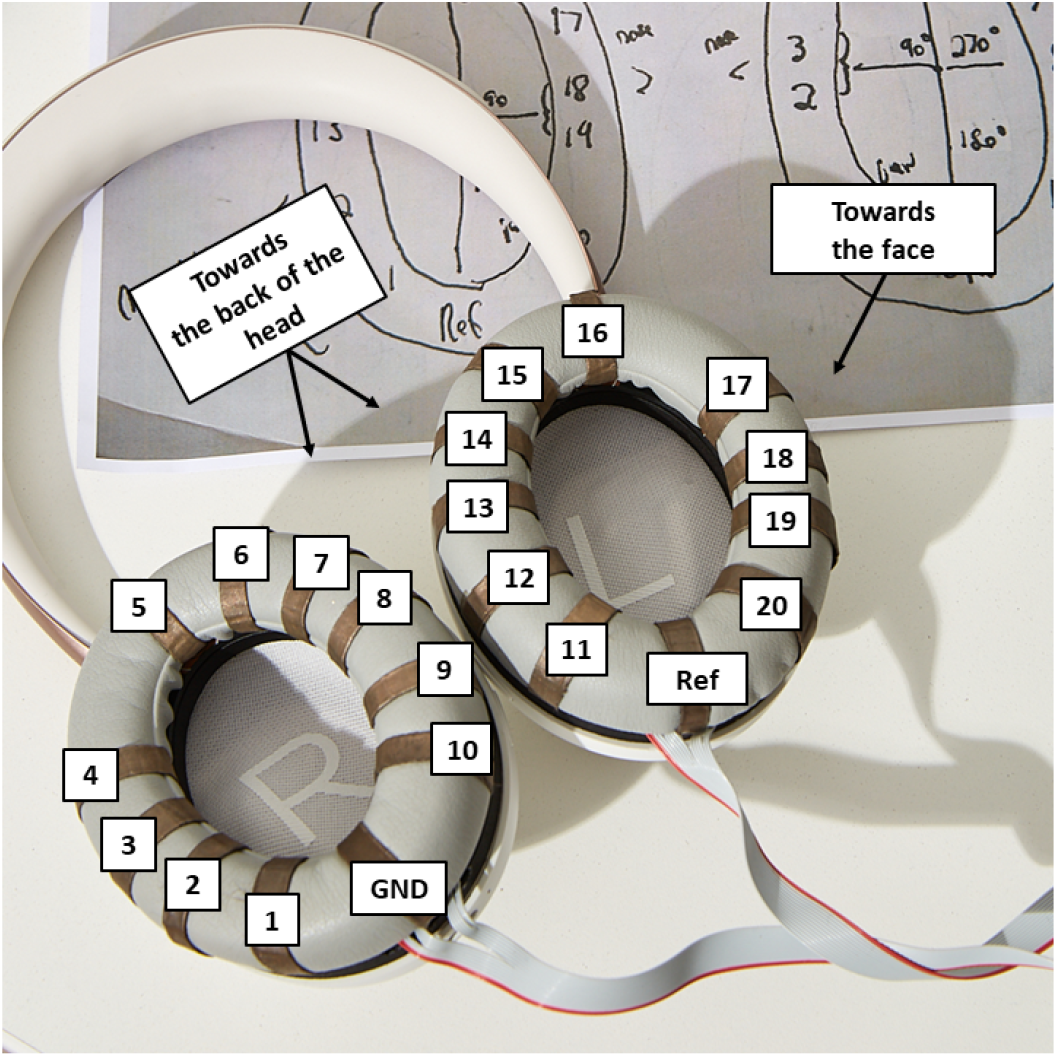
Enten headphones prototype. There are 10 electrodes on both ear pads, plus the reference (Ref) and the ground (GND) electrodes. The electrodes are dry, active, and made of a conductive mesh that wraps around the ear pads.

## 3 Validation of Enten EEG

To examine whether Enten can reliably capture neural activity, we analyzed the EEG data recorded in three different experiments with known neural responses. In parallel, we performed these experiments with a research-grade headset called Quick20 from the Cognionics company. The Quick20 is a dry electrode system that uses the standard 10-20 configuration, and has been used in other published research^2^.

The collective experiments provide preliminary validation of the Enten Headphones in a limited number of participants. If we find the expected results across all three well-known experiments, this will be early evidence that the Enten Headphones have the same response as research grade EEG headsets.

The first experiment targets the visual cortex as we record EEG data while an individual is instructed to close their eyes. In this experiment, we expect to see an increase in the alpha (8-13 Hz) activity across brain nodes. The second experiment targets the auditory cortex, in which we record EEG data in response to an auditory stimulus. In this experiment, we expect a pronounced peak in the EEG response at the same frequency as the auditory stimulus’ modulation. The third experiment corresponds to an oddball paradigm, where the participant responds to an infrequent stimulus on the screen. We expect to see a large positive potential (P300) in response. Finally, we measured the impedance of the EEG electrodes in Enten over time to ensure that good skin coupling could be achieved and that the presence of hair around the ears did not interfere with EEG contacts. Low impedance values ensure Enten captures high quality EEG signals.

### 3.1 Eyes open and closed experiment

For the first experiment, we expect the eyes-closed state to evoke more alpha oscillations compared to eye-opened state, as per the Berger effect^3^. Participants were instructed to keep their eyes open for 30 seconds, followed by 30 seconds of keeping their eyes closed. This procedure was repeated twice for both Enten headphones and Quick20. We then used power spectrogram analysis to examine changes in the alpha oscillations. Figure 2 shows the EEG spectrogram for Enten headphones and Quick20.

**Figure 2.**
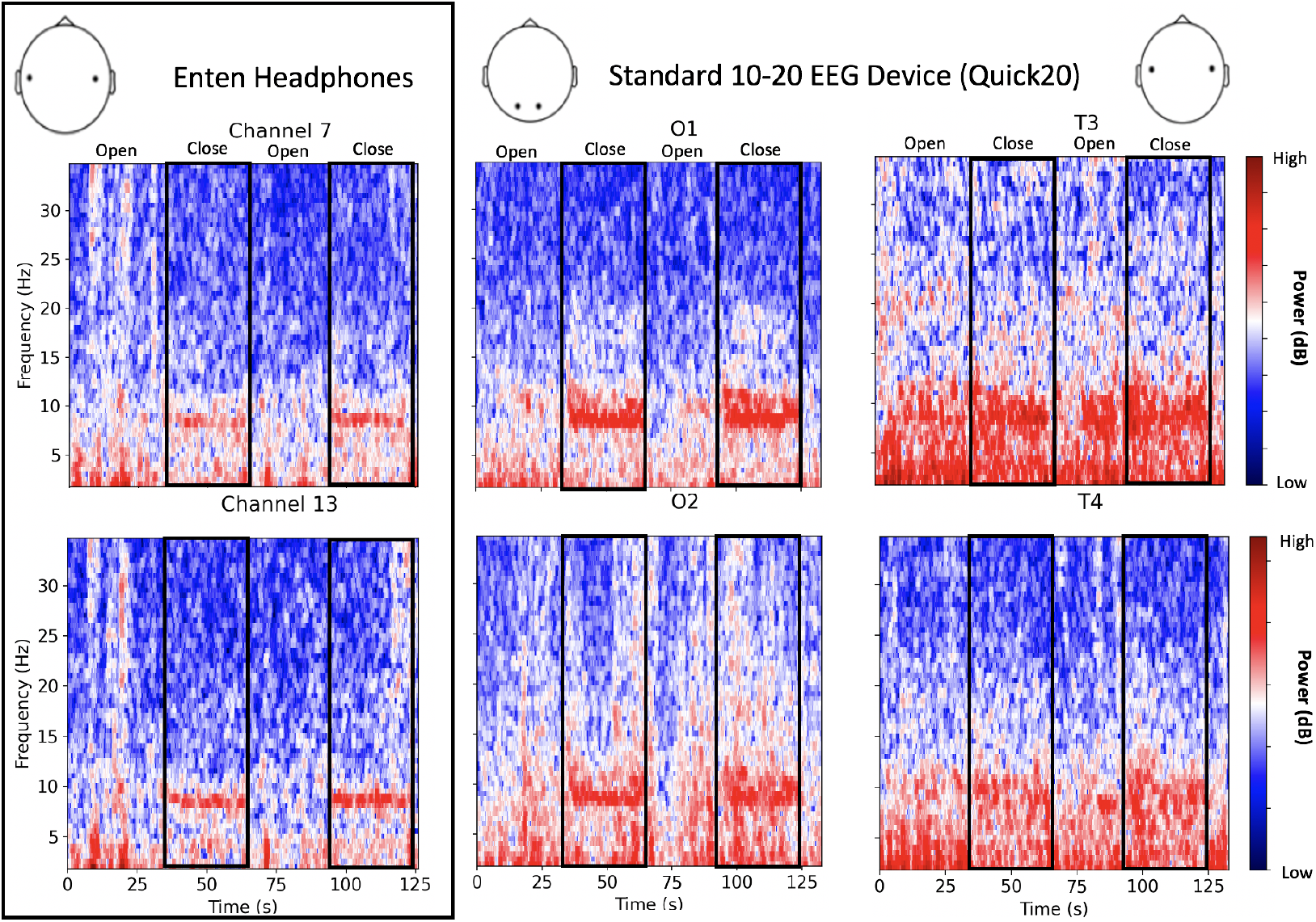
Spectrogram from the eyes opened and closed experiment. This recording is from a single participant during 2 minutes of experiment, where the eyes are open for 30 seconds followed by 30 seconds of eyes-closed state. The procedure is repeated twice, as recorded in Enten headphones (left) and Quick20 (middle and right panels correspond to the occipital and temporal channels, respectively).

Alpha waves are mostly concentrated in the visual cortex in the occipital lobe. They can also be observed in the temporal lobe. In Quick20, O1 and O2 are the electrodes located in the occipital lobe, whereas T3 and T4 are located in the temporal cortex. As seen in Figure 2, high levels of alpha oscillation - corresponding to darker red around 8 to 13 Hz - can be observed in these 4 electrodes’ locations when the participants were instructed to close their eyes. Enten headphones are able to detect these increases in Alpha oscillation, similar to Quick20.

To better quantify the difference seen in Figure 2, we also estimate power spectral densities (PSD). Figure 3 shows the PSD for Enten headphones and Quick20, which are calculated using two seconds windows. The middle and right panels of Figure 3 show the results using Quick20. We also see an increase in oscillation activity at frequencies in the beta band range (15-20 Hz). This variability in beta power is outside the scope of this experiment, but it has been reported in similar eyes open/closed experiments^4^.

**Figure 3.**
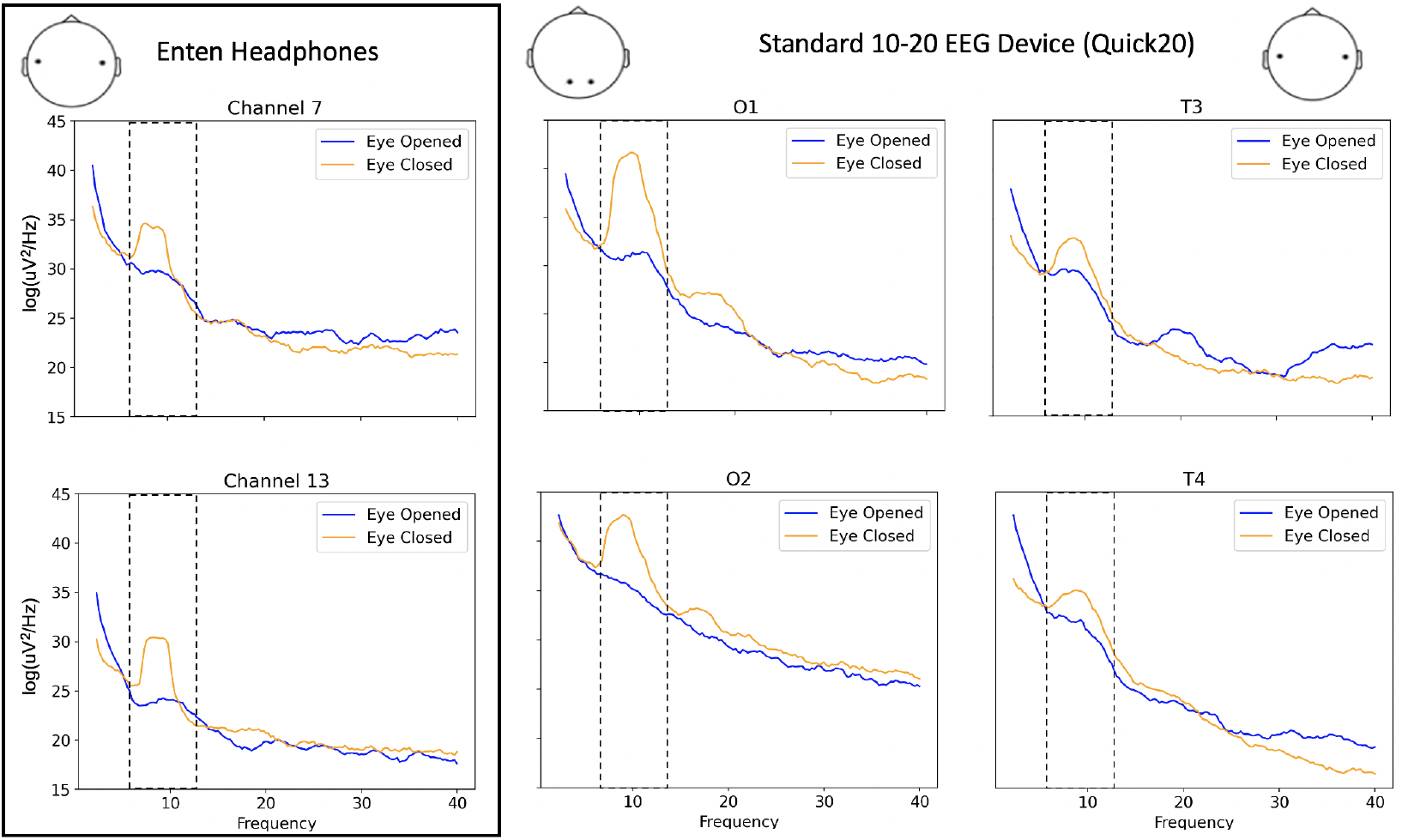
Power spectral density (PSD) for the eyes opened and closed experiment. PSD is calculated based on 4 seconds windows from the recorded data of Enten headphones (left) and Quick20 (middle and right) of the same subject. The PSD of the eyes-closed state is shown in orange, while the PSD of the eyes-opened state is shown in blue.

The left panel of Figure 3 shows the results using Enten headphones. We note that alpha power increases by about 4.0 dB and 6.0 dB for Channels 7 and 13, respectively. In contrast, with Quick 20, alpha power increases by 3.0 dB, 3.0 dB for T3 and T4, respectively. This suggests that Enten is at least on-par, or perhaps slightly better, at picking up ear-centric alpha than Quick20.

Supplementary Figure 19 expands Figure 3 across all of Enten’s electrodes and Supplementary Figure 20 further shows the comparison between Quick20 and Enten. We can see these differences between eyes opened and closed are similar to those observed for the Quick20 device, validating Enten’s EEG technology.

### 3.2 Auditory response experiment

The auditory steady-state response (ASSR) is an evoked potential elicited in response to modulated tones^5^. When a person perceives such a tone, EEG will follow the envelope of the tone. This means that if the tone was modulated at a specific frequency, we expect to see an increase in that frequency in the EEG signal, particularly in electrodes T3 and T4^6^, on the auditory cortex, as well as in electrode Cz^7^.

We played a 1000 Hz tone modulated at 37 Hz and 43 Hz^7^ via the external speakers of a desktop computer at 70% volume, for two minutes each. Figure 4 shows the PSD for the Enten headphones and the Quick20 device from two sample participants, one corresponding to each device.

**Figure 4.**
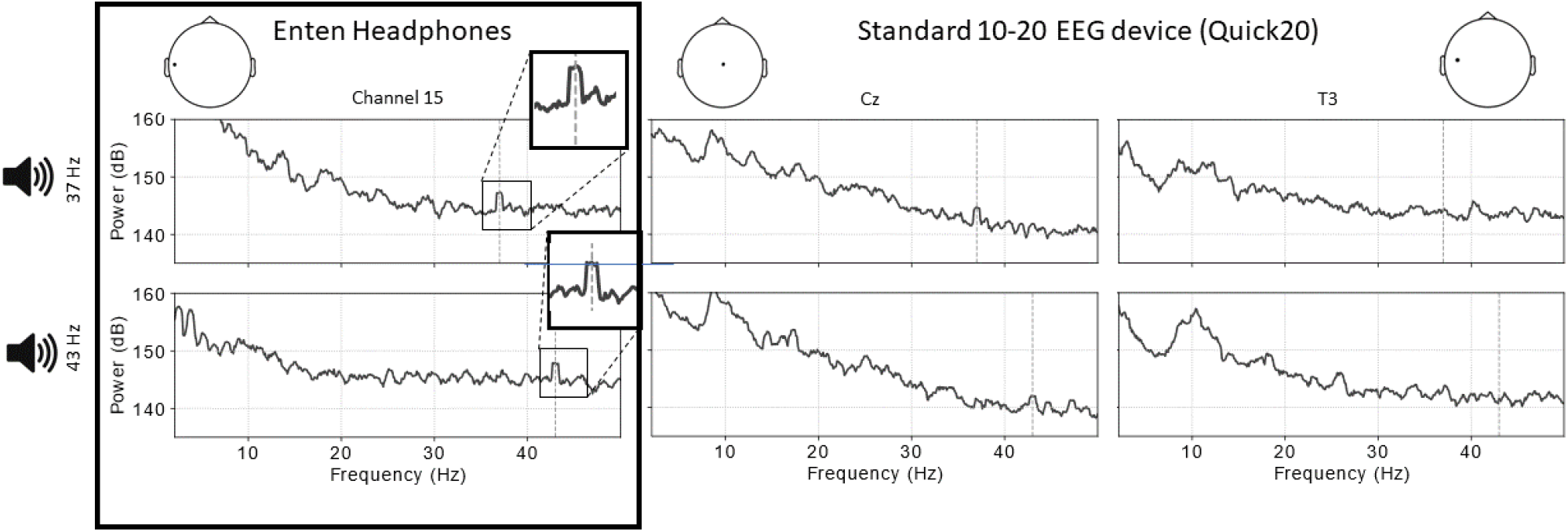
Power spectral density in audio experiment. On the left, we can clearly see the peaks in the power spectral density from Enten for a participant listening to the 37 Hz modulated tone (top) and the 43 Hz modulated tone (bottom). In the middle is the PSD using Quick20 at channel Cz. In the left and middle plots, there are clear peaks at 37 Hz and 43 Hz. Right plots show the PSD for one of the temporal channels (T3) of the Quick20. Here, the ASSR is not visible at all in either frequency

For Enten, we can see the auditory response to the tone in channels toward the top of the ear, for both frequencies. For channel 15, on the left side of the head, the peak has a height of 14 dB for 37 Hz and 14 dB for 43 Hz. For channel 6, on the right side of the head, the peak has a height of 8 dB for both 37 Hz and 43 Hz. When using the standard EEG headset, Quick20, there is a clear peak at 37 Hz and 43 Hz for channel Cz. The peak height is 14 dB for 37 Hz and 7 dB for 43 Hz. However, when observing T3 and T4, we do not see a clear auditory response with the Quick20 (for T3: 5 dB and -2 dB for 37 Hz and 43 Hz, respectively).

For both visual and auditory experiments, the Enten headphones capture the signals with a higher sensitivity compared to electrodes T3 and T4 in Quick20. This could be explained by the fact that these electrodes lie on the hair of participants, while the ones from Enten lie directly on the skin around the ears. Even though the electrodes on Quick20 are designed as small spikes, the surface area in contact with the scalp is smaller than the electrodes that lie directly on the skin. The scalp also has different characteristics compared to the exposed skin, which increases the impedance of the connection.

### 3.3 P300 experiment

The P300 is a well known potential evoked in response to infrequent or target stimuli. When people are exposed to a stream of stimuli, they will produce a positive deflection around 300 ms after stimulus presentation (P300) for stimuli that either are infrequent or require a response^8^.

In this experiment, one participant was presented with 4 words, and was instructed to press a button only when the word “BLUE” appeared on the screen. The experiment had 200 trials of 1.2 seconds with an inter-trial time interval of 2-3 seconds.

The words “BLUE” appeared randomly 25% of the time. This experiment was performed in both Quick20 and Enten. The results are presented in Figure 5. On the left panel, we show the response averaged across trials for Enten over channel 6. The difference between the P300 and the baseline is 4 uV. On the middle panel we show the results for Quick20 over the occipital cortex (O1); besides the P300 we also observe the visual evoked potentials, such as the P100. The difference between the P300 and the baseline is 3.5 uV. Finally, on the right panel, we show the results for Quick20 over the right temporal cortex (T4), where we see a difference of 5 uV between the peak of the P300 and the baseline.

**Figure 5.**
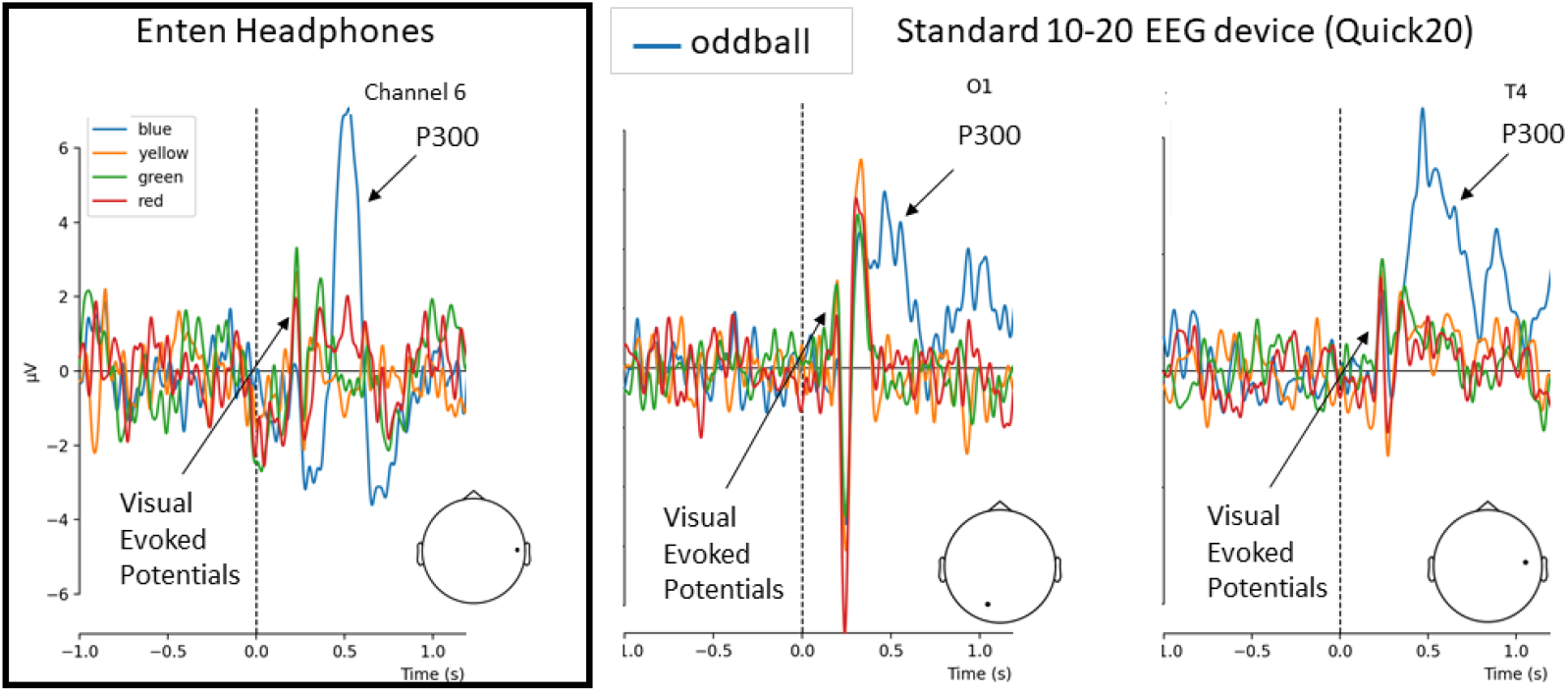
Evoked response from Oddball experiment for Quick20 and Enten. The participant was placed in front of a computer screen with different color words appearing on the screen and was instructed to press a key every time the word “BLUE” appeared on the screen, and to ignore all other words. This experiment was performed twice, once on the Quick20, and once on Enten.

In conclusion, we saw a P300 of similar amplitude for both Quick20 and Enten. The visual evoked potentials are clearly present in the occipital channels of Quick20, and barely present for both the temporal channels of Quick 20 and for Enten.

### 3.4 Impedances

As an additional test to ensure signal quality, we evaluated channel impedances over time. Impedance provides a measure of the channel’s degree of coupling to the skin, with higher values indicating poorer contact and therefore poorer reading of neural signals. We found that impedances reached desirable levels (<300 kΩ) after a settling time that was on the order of minutes. One of these test runs is shown in Supplementary Figure 21, during which over half of the channels settled to a low impedance (<100 kΩ) after 10 minutes, and 17 channels had reached this value after 16 minutes of wear. An impedance threshold of 100 kΩ has been used in several EEG studies (e.g., Mathewson et al.^9^).

## 4 Behavioral changes in response to cognitive demands and distractions

It is often argued that focus is both dependent on the changes in the external environment and a person’s intrinsic will to concentrate^10^. Focus is the ability of a person to concentrate on a task without reacting to external distractions or internal changes in motivation. To build our focus analytics platform and to validate its performance, we have designed a cohort of cognitive experiments that let us investigate how the brain and behavior are influenced by changes in focus and distractions.

In our experiments, we use variations of the Stroop experiment^11^ by embedding different visual backgrounds and audio distractions in the context of the Stroop experiments. These variations let us change the level of focus in a controlled manner. We argue these variations help us to better simulate changes in focus level in a replica of real-world settings. This viewpoint is aligned with more recent research efforts that use the Stroop task and its variants, like affective and emotional stroop, to study focus and other domains of cognitive functions^12,13^. In the following subsections, we present a brief explanation of the experiments we conducted at Neurable along with analysis of behavioral outcomes.

### 4.1 Experiment 1 - Distraction Stroop Task

During each trial of this experiment, colored words (e.g., Red or Yellow) appeared on the screen. The participants were asked to respond to the color of the words and ignore their meaning by pressing four keys on the keyboard – “D”, “F”, “J”, “K” – mapped to - “Red”, “Green”, “Blue” and “Yellow” - colors, respectively. Trials in the Stroop task are categorized into congruent, when the text content matches the text color (e.g., Red) and incongruent, when the text content does not match the text color (e.g., Red). The incongruent case is counter intuitive and more difficult for the participants to provide a correct response. Behavioral outcomes (reaction time, accuracy) drops in incongruent trials (lower accuracy, higher response time) which is known as the Stroop effect^11^.

To study the Stroop effect in more unconstrained settings, participants were required to perform the task while being bombarded with additional sensory information in the form of audio and video. We hypothesize that additional sensory information will let the Stroop effect to be more salient and represent distractions present in the daily life settings. The background distractors embedded in the experiment were: a calm river scene, a thrilling roller coaster point of view scene, a soothing beach scene, and a noisy marketplace scene. These videos provide different levels of distraction which interfere with a person’s sustained focus to respond to the Stroop trials.

Participants performed 4 blocks of the Stroop task, each block containing 64 trials. There were equal numbers of congruent and incongruent trials. Each block corresponded to a different background. Participants had 1.2 seconds per trial to respond to the task. If there was no answer in this interval the trial was marked as missed and the experiment went to the next trial. Both behavioral and neural data of 64 unique participants, totalling 80 sessions, was collected using Enten.

Figure 6 shows analysis of behavioral data across participants. The average reaction time is longer, average accuracy is lower and proportion of missed trials were higher during all blocks with incongruent trials. These results align with the Stroop effect. Whilst the stroop effect is present in each block, we were able to change the dynamics of the effect from one block to the other. For instance, during the roller coaster background (second row) the participants had the highest accuracy with the least number of missed trials. During the marketplace background (bottom row), the participants had the worst accuracy and highest proportion of missed trials, indicating that it was the most distracting background. Significant changes in neural activity captured during the experiment, which are presented in the next sections, also show promising effects brought on by changes in background. In particular, for the marketplace background, there is a consistent drop in the alpha band power during the incongruent trials and the magnitude of the drop is the highest across backgrounds (Figure 10, Experiment 1A. Marketplace). Per the result extracted in our Stroop distraction experiment, we decided to examine in what conditions the changes in the behavior would be more salient. Our hypothesis was that the distracting marketplace background combined with a higher percentage of incongruent trials would make the stroop task more challenging, whereas no background sensory information - like plain grey - with congruent trials will be the least challenging setting. As a result, we designed Experiment 1B. To drive the behaviour towards two extremes and capture larger deviations across conditions, we divided the experiment into two blocks: in the first block, participants undertook a stroop task with all congruent trials in a plain background (Easy Stroop), and the second block, participants respond to Stroop task with a high percentage of incongruent trials (75% incongruent), while being distracted by the marketplace background (Hard Stroop). Behavioral and neural data of 21 unique participants was collected using Enten for Experiment 1B. Figure 7 shows behavioral data across these participants and the easy and hard blocks. The expected gap between reaction time in the congruent and incongruent blocks went from 70ms in experiment 1A to 140 ms between the easy and hard blocks in experiment 1B. This result from behavioural data aligns well with our hypothesis in successfully modulating the difficulty of the task and magnifying its effect on the statistical properties of corresponding neural correlates (e.g Alpha power band) which can be used as a possible neural marker in our proxy of focus, which will be discussed in section 5.

**Figure 6.**
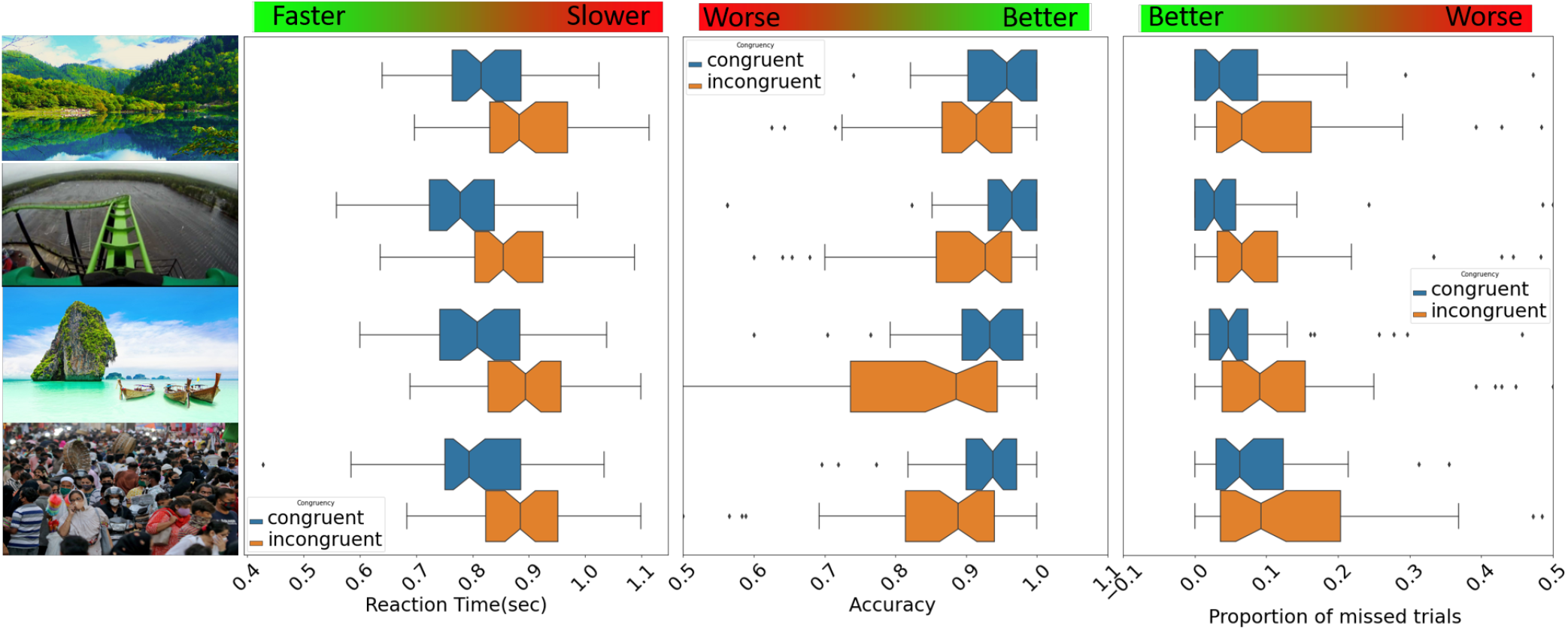
Stroop Task and Behavior Analysis. The leftmost column represents the four blocks of the experiment background, namely a calm river, a roller coaster, a calm beach and a noisy marketplace. Mean reaction time (second column), mean accuracy (third column) and proportion of missed trials (fourth column) were calculated per subject, then averaged across the population as a function of congruent and incongruent trials. Incongruent trials were characterized by longer reaction times, lower accuracy, and higher proportion of missed trials as compared to congruent.

**Figure 7.**
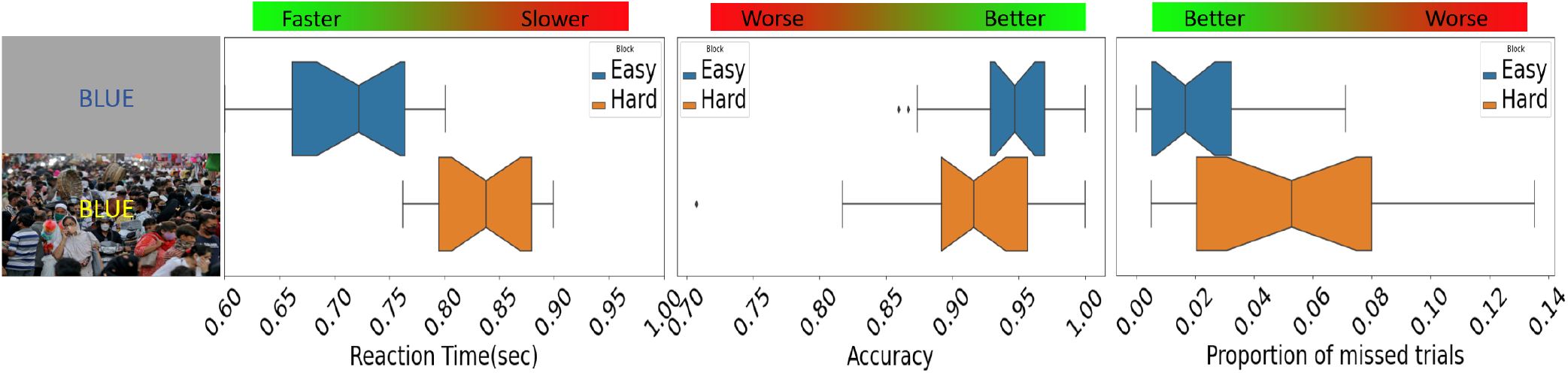
Experiment 1B and Behavior Analysis. Two blocks of the experiment background (first column), Easy, and Hard. Mean reaction time (second column), mean accuracy (third column) and proportion of missed trials (fourth column) were calculated per subject, then averaged across the population as a function of easy and hard blocks. Hard blocks were characterized by longer reaction times, lower accuracy, and higher proportion of missed trials as compared to easy.

### 4.2 Experiment 2 - Interruption by notification

In Experiment 1, task difficulty was modulated with different ratios of congruent and incongruent trials in the blocks, as well as with video backgrounds presenting different levels of complexity in sensory information. In Experiment 2, we tested how our focus prediction algorithm, which we present in section 5, responded to known distractions, and whether this effect was consistent across the population. It is widely established that push notifications will decrease focus level^14^. As a result, we embedded this type of distraction in our experiments and examined how our focus metrics respond to it.

Experiment 2 had three variations. The first variation (Experiment 2A) was a replica of Experiment 1B, with the addition of notification sounds that appeared randomly during some blocks of the experiment. When participants heard the notification sound, they stopped performing the main task (Stroop) and responded to a chatbot (cleverbot.com). They would then resume the main task. The left panel of Figure 8 depicts the experiment design for Experiment 2A. Our behavioral analysis shows that, on average, participants presented slower reaction times and were less accurate during blocks with distractions compared to blocks with no distractions. This is shown in Figure 9.

**Figure 8.**
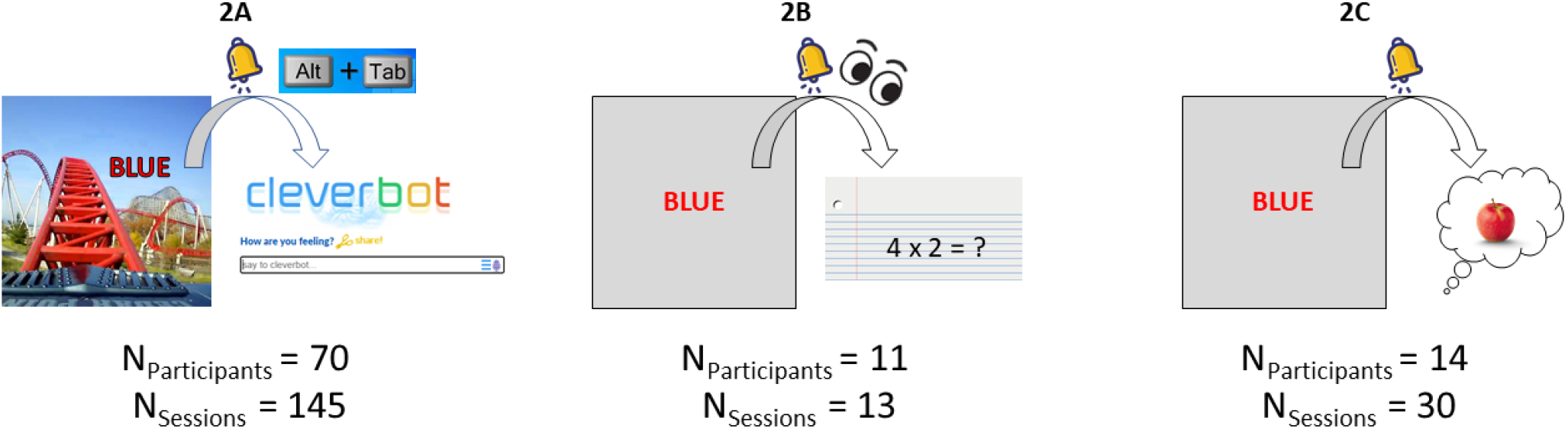
Summary of variations of Experiment 2. In Experiment 2A, participants performed the Stroop task without distractions during some blocks and with distractions in other blocks. The distractions consisted of a notification sound that was played at random times. When participants heard this sound, they had to switch windows, respond to a chatbot, and then quickly return to the Stroop task. In experiment 2B, every time participants heard the notification, they had to solve a math problem silently shown on a piece of paper below the monitor. In Experiment 2C, when participants heard the notification, they had to recall an object that was the same color as the word on the screen. Below each experiment is the number of participants and valid sessions for each experiment.

**Figure 9.**
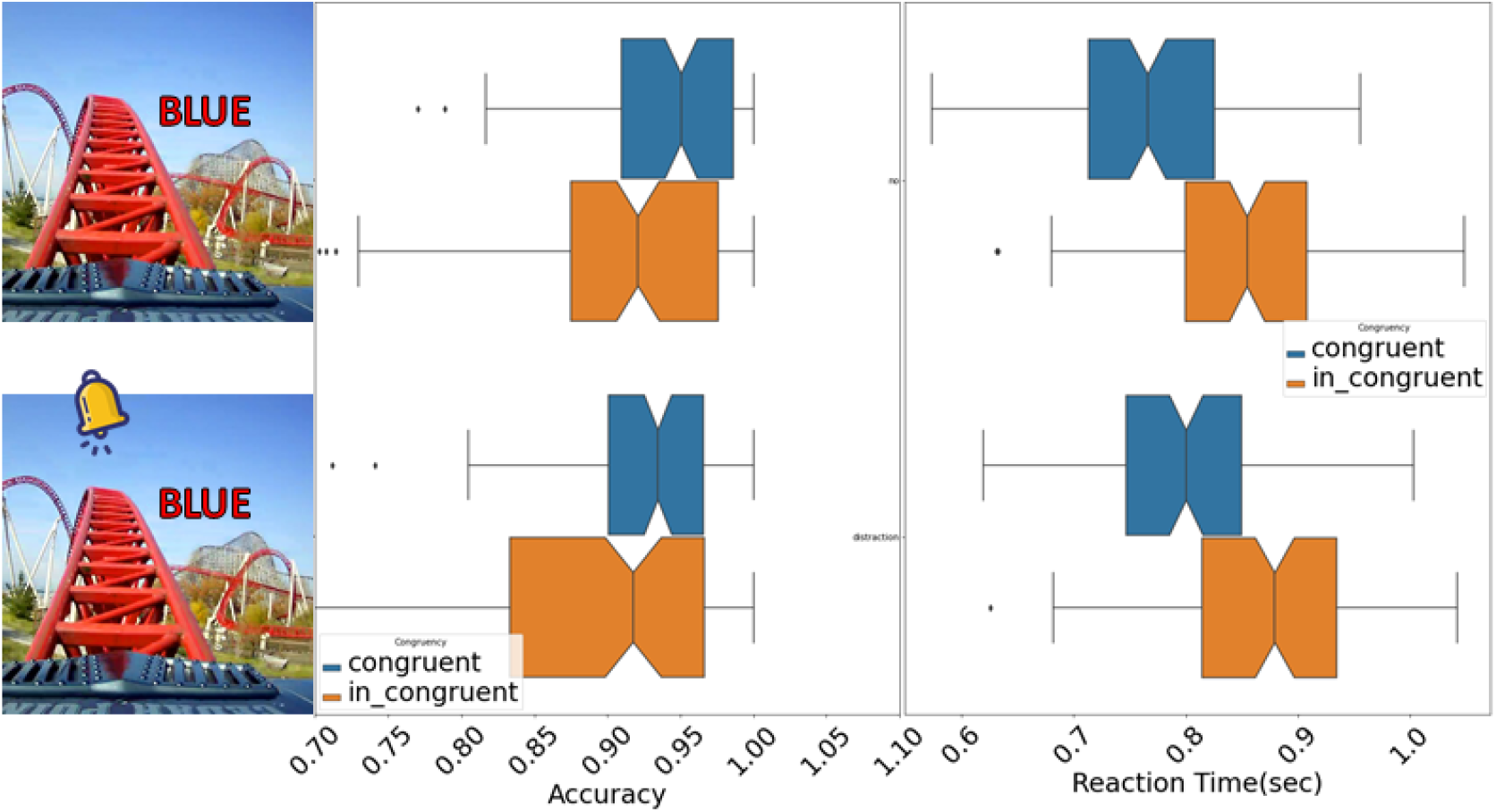
Behavioral responses during Experiment 2A. Accuracy (middle) and reaction time (right) grouped by whether the trials were in a block without distractions (top row) or with distractions (bottom row). Only trials where there was a response within the allotted time (1.2s) were included in this analysis.

Working with Enten in an unconstrained setting will bring up the question of whether our focus algorithm is primarily driven by neural activity, or whether it is leveraging confounding signals to predict focus, such as EMG or motion artifacts. To ensure that this was not the case, we adjusted Experiment 2A to minimize potential noise sources. While the distraction is present in all variations, we adjusted tasks performed in response to the notification to control for eye and body movement. To reduce the number of tested variables, in Experiments 2B and 2C, we kept the background and the level of stroop congruency consistent throughout the experiment. In Experiment 2B, participants solved a mental math problem presented on a sheet of paper below the monitor every time they heard the notification sound (Figure 8, center). This design reduced body movements during distractions, but it did not control for sources of artifact caused by eye movements. In Experiment 2C, participants were asked to keep their eyes on a fixation cross. When participants heard a notification, they were asked to recall an object of the same color as the word appearing on the screen (Figure 8, right). In both Experiments 2B and 2C, the stroop congruency level and background were held constant throughout the experiment. Our behavioral analysis shows results that are consistent with Experiment 2A. Later in Section 7, we will present evidence that the EEG artifacts were successfully reduced in Experiments 2B and 2C compared to Experiment 2A.

## 5 Neural activity changes in response to distraction

As described earlier, a distraction is any sensory input (external or internally generated) that moves our focus away from the attended task. It has been shown that maintaining a high neural activity in the alpha band distributed across the brain helps us filter out and ignore distractions^15^. When alpha activity decreases, it indicates a loss of focus. Activity in other frequency bands, such as beta (13-30 Hz), is also associated with attention^16^.

To show changes in neural activity in response to distractions, we considered incongruence in Stroop trials as distractions. We compared power in the alpha band in congruent and incongruent trials in different blocks of Experiment 1A. We normalized the power by subtracting the average power in the alpha band in the 200 ms before the beginning of the trial. Our analysis shows that alpha power is consistently lower in incongruent trials when compared to congruent ones (Figure 10). This effect was particularly strong with the marketplace background, which proved to be the most distracting scene. A similar reduction of neural activity in incongruent trials was observed in the low beta band (13-20 Hz). We then analyzed the data in Experiment 1B, which was divided into easy and hard stroop blocks. Hard blocks utilized the marketplace as background and had a higher number of incongruent trials. Hence, we hypothesized that the power in alpha (and low beta) frequencies should be attenuated during these hard blocks. We normalized each block with the preceding 500 ms, and then averaged all blocks together (Figure 11). The results confirmed our hypothesis, showing that alpha and low beta power are consistently lower in the hard block than in the easy block. Taken together, these results demonstrate that our experiments successfully modulated focus and that incongruent trials cause a loss of focus that is detectable in the participant’s brain activity.

**Figure 10.**
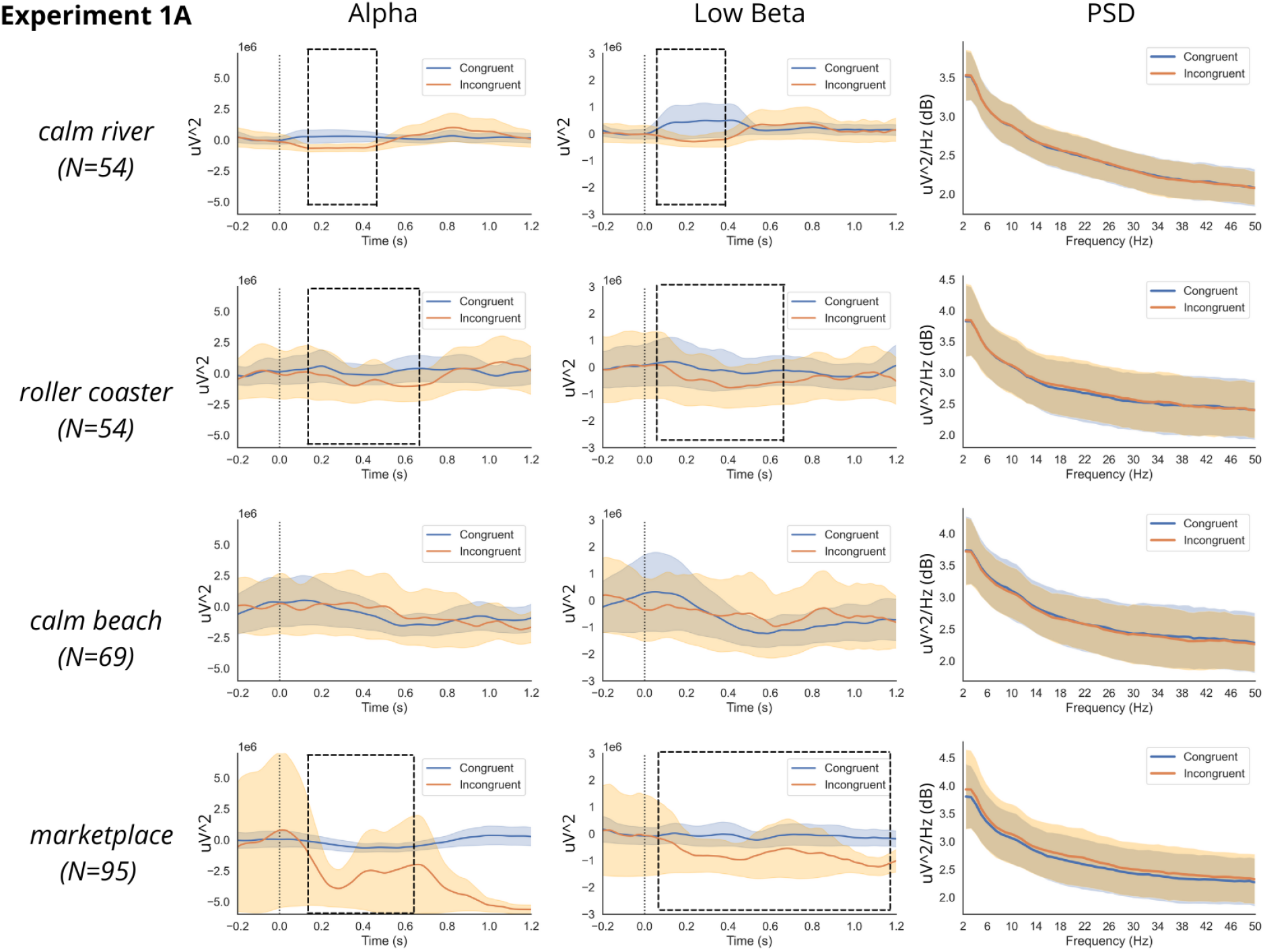
Analysis of neural activity during Experiments 1A. Temporal dynamics of power in alpha and low beta frequency bands in congruent and incongruent trials for channel 6. Incongruent trials are accompanied by a reduction in alpha and low beta power, which we consider a proxy for loss of focus (see black rectangles). These differences were not due to gross EMG artifacts, as the PSD was similar between congruent/incongruent trials and easy/hard blocks (third column). Note that these changes are too small to be seen in the PSD plots. For all plots, shaded areas represent standard error of the mean across participants. Some participants performed Experiment 1A with only a subset of backgrounds due to time limitations, hence the different N across backgrounds.

**Figure 11.**
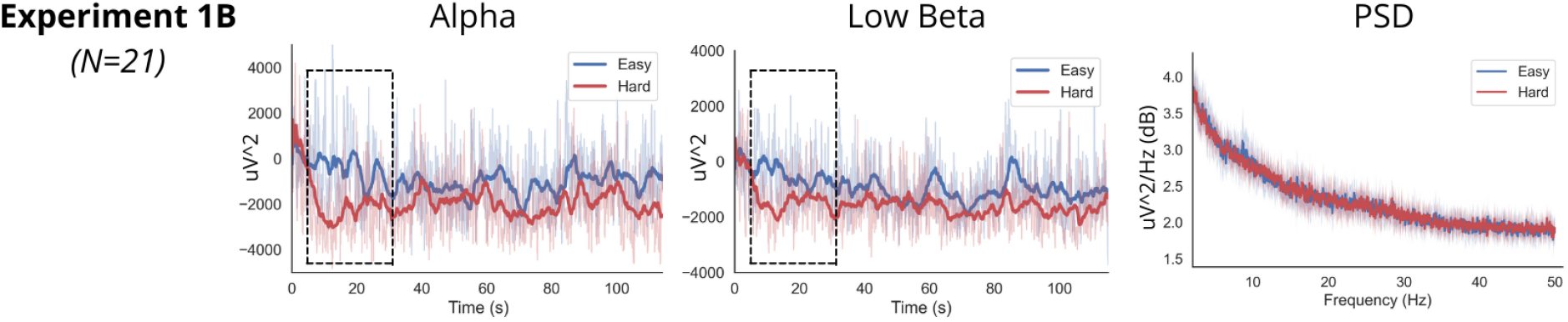
Analysis of neural activity during Experiments 1B. Temporal dynamics of the power in alpha and low beta frequency bands in easy and hard blocks. Thick lines represent moving averages of thin lines (normalized alpha and low beta power), with a window size of 1.2 seconds. Hard blocks (characterized by more incongruent trials and marketplace background) cause a perceivable loss of focus and a consequent reduction in alpha and low beta power (black rectangles). These differences are not due to gross EMG artifacts, as the PSD was similar between blocks (third column). Note that these changes are too small to be seen in the PSD plots.

Collective results in Experiment 1 show that distractions cause changes in the neural activity that Enten captures. The strongest neural patterns associated with focus are in the alpha band and are robust to overall increase in power caused by potential EMG activity. Distractions also cause changes in power in other frequency bands, such as low beta (see Figure 10,11), which could be incorporated in estimation of focus, but are not the aim of this white paper.

## 6 Neurable’s ML model detects distractions in EEG

After validating the presence of neural signatures of focus and the impact that distractions have on observed neural dynamics across our experiments, we explored ML techniques to translate changes in neural activity into measurable estimates of focus. In the previous section, we have seen that, in Experiment 1B, the focus level of participants was lower in hard blocks than in easy blocks of Stroop. This was confirmed by behavioral data, which showed lower accuracy and longer reaction times in hard blocks. Hence, we trained a support vector machine (SVM) classifier to predict whether the user was performing an easy task, which is the proxy of a condition in which it is easier to focus, or a hard task, a proxy of the condition in which it is hard to focus, using the data from Experiment 1B. This classifier used the power in the EEG alpha band at different electrodes as features. To account for overall differences in power, we normalized each feature by dividing its value by the total power in that channel. We also scaled features by subtracting the median and dividing by the interquartile range (i.e., difference between first and third quartiles), computed on the whole dataset. This scaling technique ensures that large outliers are not taken into account in the scaling procedure, as opposed to traditional z-scoring. This step makes the data from one session comparable to the data from another session, irrespective of any difference in magnitude between features due, for example, to different impedance levels across days. We used the probability that the participant is engaged in an easy task, as estimated by the SVM, as a proxy for your focus. Specifically, a higher level of alpha characterizes easy tasks and corresponds to higher focus. To test our model in realistic settings, we hypothesized that distractions of Experiments 2A-C increase the difficulty of the task and make the user lose focus, hence being analogous to hard blocks in Experiment 1B. This was motivated by our aim to build a model for decoding focus independent of the particular task at hand or distraction type (i.e., incongruent trials or sound). Hence, we tested our model on data from Experiments 2A-C, where participants were receiving auditory distractions.

First, we used our algorithm to estimate the focus score over time for a single user undertaking Experiments 2A-C over different days. The focus score ranges from 0 to 100, where larger numbers mean a higher level of focus (Figure 12). Each value of the focus score was derived by the model from the neural activity in the alpha band. To capture reasonable alpha changes in the context of focus dynamics while controlling spurious noise in our model estimates, we smoothed the focus prediction using a moving average of the most recent 10 focus predictions. Our hypothesis was that the focus score should be high when there are no distractions, and should drop as participants get distracted or lose focus. As hypothesized, the focus was overall much higher in experimental blocks without distractions (denoted by green lines on top) than in blocks with distractions (magenta lines on top). Moreover, a distraction event caused an immediate drop in focus.

**Figure 12.**
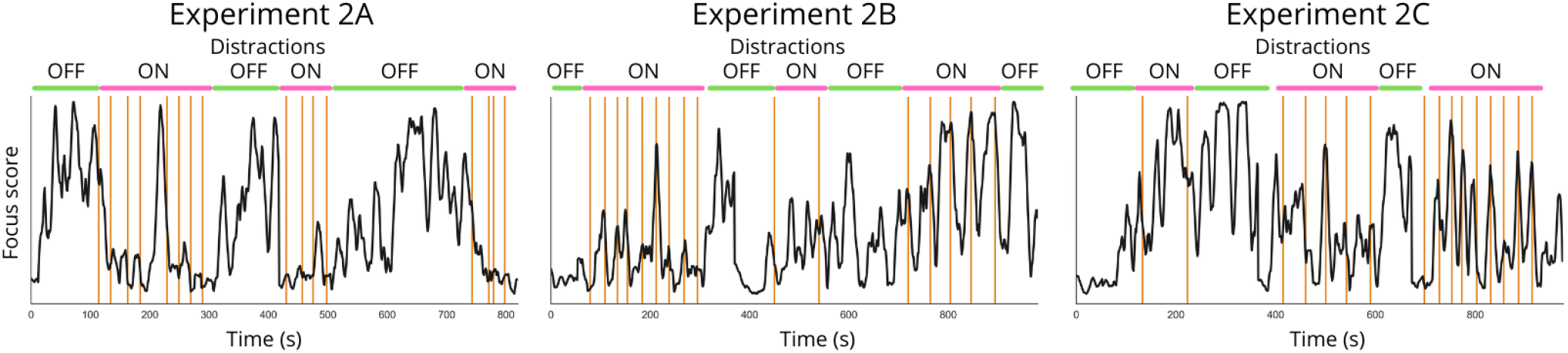
Focus score accurately represents user focus across different experiments. Focus score estimated by our algorithm for one participant undertaking Experiments 2A-C on different days. Each column represents data from a different experiment. Vertical orange lines indicate times of the distraction sound. As hypothesized, the focus score is much higher in blocks without distractions than in blocks with distractions. Distraction events cause an immediate loss of focus, and participants take time to regain their focus over the course of an experiment.

While visual inspection of temporal dynamics of focus seems to reveal a positive and consistent correlation with the experimental blocks, we developed a specific metric (QA score) to assess how accurate our algorithm was in detecting the occurrence of the distractions embedded in our experiments. Specifically, we looked at how fast the focus score changed from one second to the next. If the first-order derivative of the focus score was below a pre-set negative threshold (i.e., drop in attention) and the focus score was below a pre-set threshold (i.e., attention is low enough), we considered this a distraction detected by the model. These constraints limited the number of false positives, allowing for natural oscillations of focus that happen over time. In our experiment, we knew at which time points the distractions were issued and, so, we could compute the sensitivity and specificity of our algorithm. The QA score was therefore computed as the average between the sensitivity and the specificity of the model in detecting distractions over the course of the experiments:

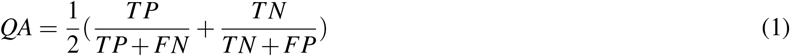

where TP are the number of true positives (i.e., correctly identified distractions), TN are the true negatives (i.e., correctly identified trials without distractions), FP are the false positives (i.e., the model detected a distraction when none was issued), and FN are the false negatives (i.e., the model did not detect a distraction that was instead present).

Figure 13 shows an example of our algorithm estimating focus for a representative participant. To easily visualize the slow variability of focus over time, we smoothed the predictions made by our model (thin grey line in Figure 13A) using a moving average with a 10 sample window length (thick black line in Figure 13A). The corresponding derivative of the focus score, used by our algorithm to detect distractions, is shown in Figure 13B. We can see a drop in the focus score in 15 out of 16 distractions administered, with very few false positives. The only distraction not correctly detected appears shortly after another distraction, suggesting that the brain did not have enough time to regain focus after the previous distraction and, as a result, it did not show a drop in focus.

**Figure 13.**
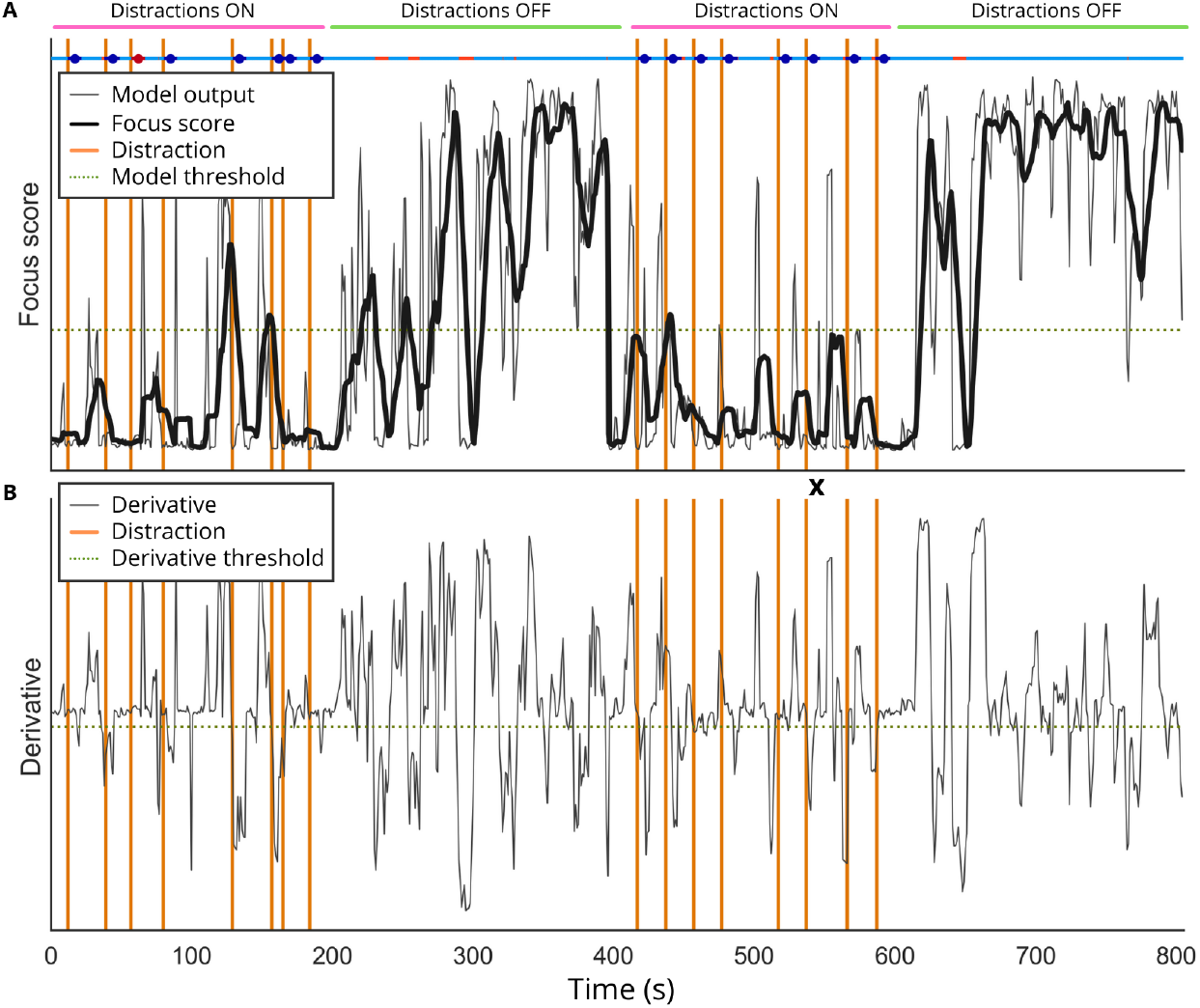
Algorithm for focus score estimation in Experiment 2A. (A) Raw model’s output (thin grey line) and smoothed focus score (thick black line) at each time point of the experiment, and (B) first-order derivative of the raw model’s output for a representative participant. Vertical orange lines indicate distraction onset. Dots on top represent correct (blue) and incorrect (orange) detections of distractions. For example, the distraction marked with a cross clearly shows a drop in the raw model’s prediction on top, and a negative derivative on the bottom. As a result, it is correctly identified by our distraction detector.

To validate the generalizability of our model across participants, we calculated the QA score for all participants undertaking Experiments 2A-C. The results are shown in Table 1, demonstrating the generalizability of our model across sessions, participants and experiments.

**Table 1.** Performance of the model detecting distractions for Experiments 2A-C across all participants as assessed by the QA score (mean ± standard deviation). Each experiment lasted approximately 15 minutes, and included between 16 and 19 distractions

| Experiment | Experiment 2A | Experiment 2B | Experiment 2C |
| --- | --- | --- | --- |
| QA score | 81.0 ± 5.0% | 77.4 ± 6.2% | 77.2 ± 5.7% |
| # of sessions | 145 | 13 | 30 |
| # of unique participants | 70 | 11 | 14 |

## 7 Algorithm robustness to artifacts

A critical feature of decoding algorithms based on neural activity is their robustness to artifacts, for example, those generated by body movements, jaw movements, and eye movements. Such movements could potentially be correlated with the behavioral task and, therefore, could potentially be used by our algorithm in lieu of neural data to predict distractions. In the previous sections, we have shown that our preprocessing pipeline allows us to detect neural activity that is independent from body movements, such as power increases in EEG alpha waves during eyes closed. We specifically designed Experiments 2A-C to check the resiliency of our algorithm for focus estimation to confounding factors, such as body and eye movements.

### 7.1 Single subject longitudinal analysis

We first examined a single subject performing Experiments 2A-C across different days. None of the three experiments show broadband increases in PSD after distraction, an effect usually associated with EMG artifacts (Figure 14A). The accelerometer data showed significantly more movements after distraction in Experiment 2A, but not in Experiments 2B-C (Figure 14B). However, for all three experiments, alpha features and the focus score predictions by our algorithm were significantly lower after distraction (Figure 14C). This suggests that, for this subject, EMG artifacts did not impact our model’s predictions. Clear alpha peaks are visible during all experiments (Figure 14A). Additional peaks are present in low beta before but not after distractions, suggesting that beta inhibition could also help our estimation of the focus^16^, a venture that Neurable will explore in future research.

**Figure 14.**
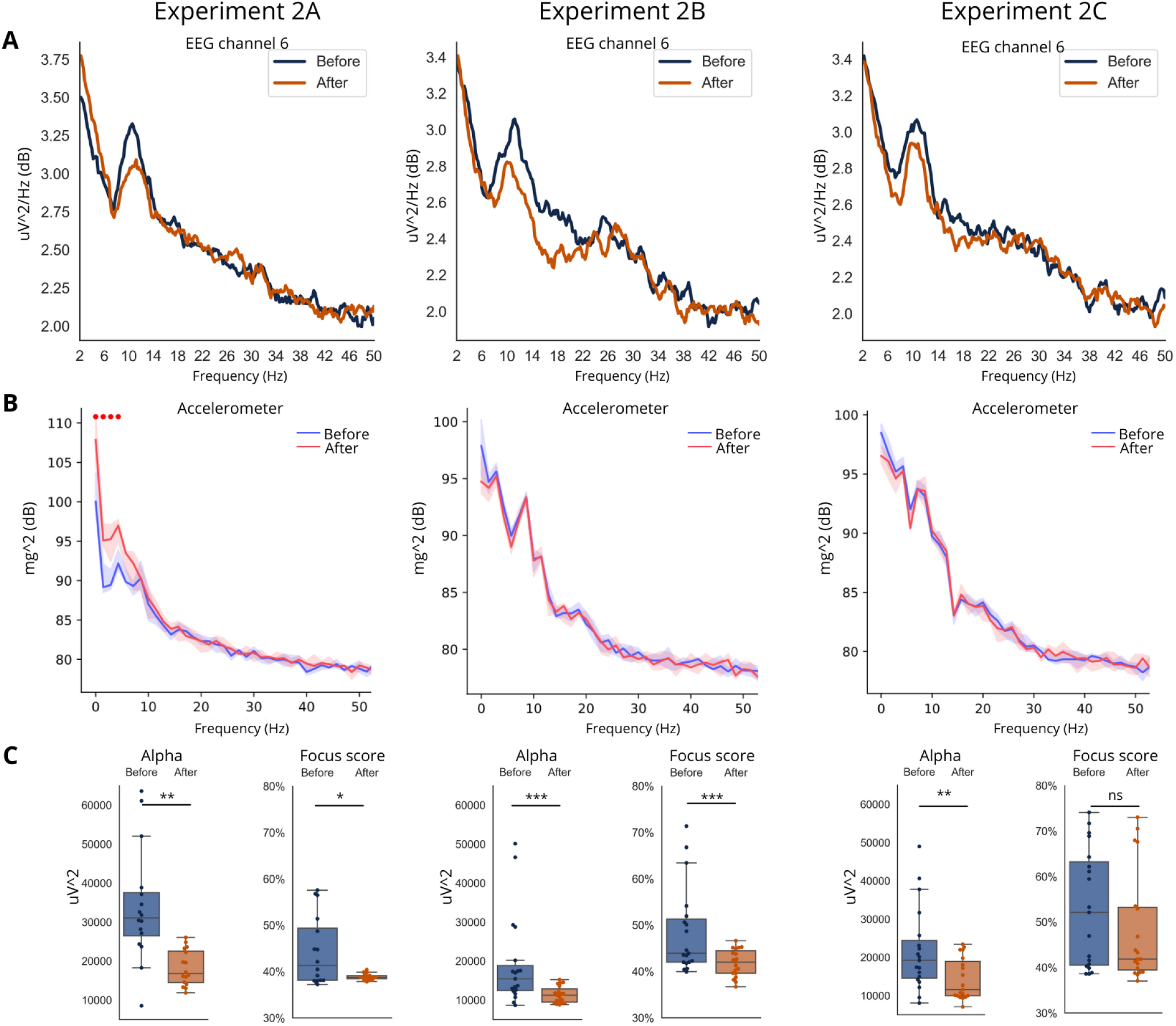
Focus algorithm is resilient to EMG artifacts at the single subject level. (A) Average PSD in log scale at channel 6, (B) PSD of the accelerometer channel, and (C) average power in alpha and model’s predictions at channel 6 before and after distractions. Each column represents data from a different experiment (Experiments 2A-C). The accelerometer data show significantly more movements after distraction in Experiment 2A (see red dots on top, p<0.001 by Wilcoxon signed-rank test), but not in Experiments 2B-C, confirming our choice of the experiment variations. Nevertheless, all experiments show clear peaks in alpha and low beta, attenuated after distraction, and comparable performance of our algorithm, suggesting its robustness to EMG activity generating artifacts. Significance of alpha and focus score was tested using Wilcoxon signed-rank test with alpha=0.01. In box plots, * corresponds to p<0.05, ** to p<0.01, *** to p<0.001, ns to p*≥*0.05.

### 7.2 Group analysis shows motion artifacts do not drive focus predictions in Experiments 2B and 2C

To further verify the robustness of our algorithm to EMG and other noise sources, we repeated the above analyses at the group level (see also Table 1, which shows that focus was robustly reported across subjects). For our group level analysis, we pooled the distraction events from all sessions together and calculated median PSDs and statistics. If the model’s performance was derived by EMG, we hypothesized that this would manifest as increases in broadband EMG and accelerometer activation following distractions. Similarly, EOG would likely show up as increased low frequency activity.

Our group level analysis of experiment 2A is shown in Figure 15 (left). We found that there were significant increases in EEG PSD following distractions. This was mostly a broadband increase, with the exception of alpha and low-beta frequencies (Figure 15, top left). Likewise, accelerometer activity increased (Figure 15, middle left). This suggests the presence of EMG, motion artifacts, and possible EOG contamination in these data. These increases were almost completely abolished during experiments 2B and 2C. Alpha, on the other hand, was significantly attenuated after distractions in experiments 2B and 2C. While there was slight attenuation of the alpha peak present in experiment 2A, this did not reach significance and was likely obfuscated by the aforementioned broadband increase in PSD. The fact that alpha is attenuated in the absence of any changes in accelerometer activity in experiments 2B and 2C suggests that, in these cases, the model is being driven by alpha EEG and not EMG or EOG.

**Figure 15.**
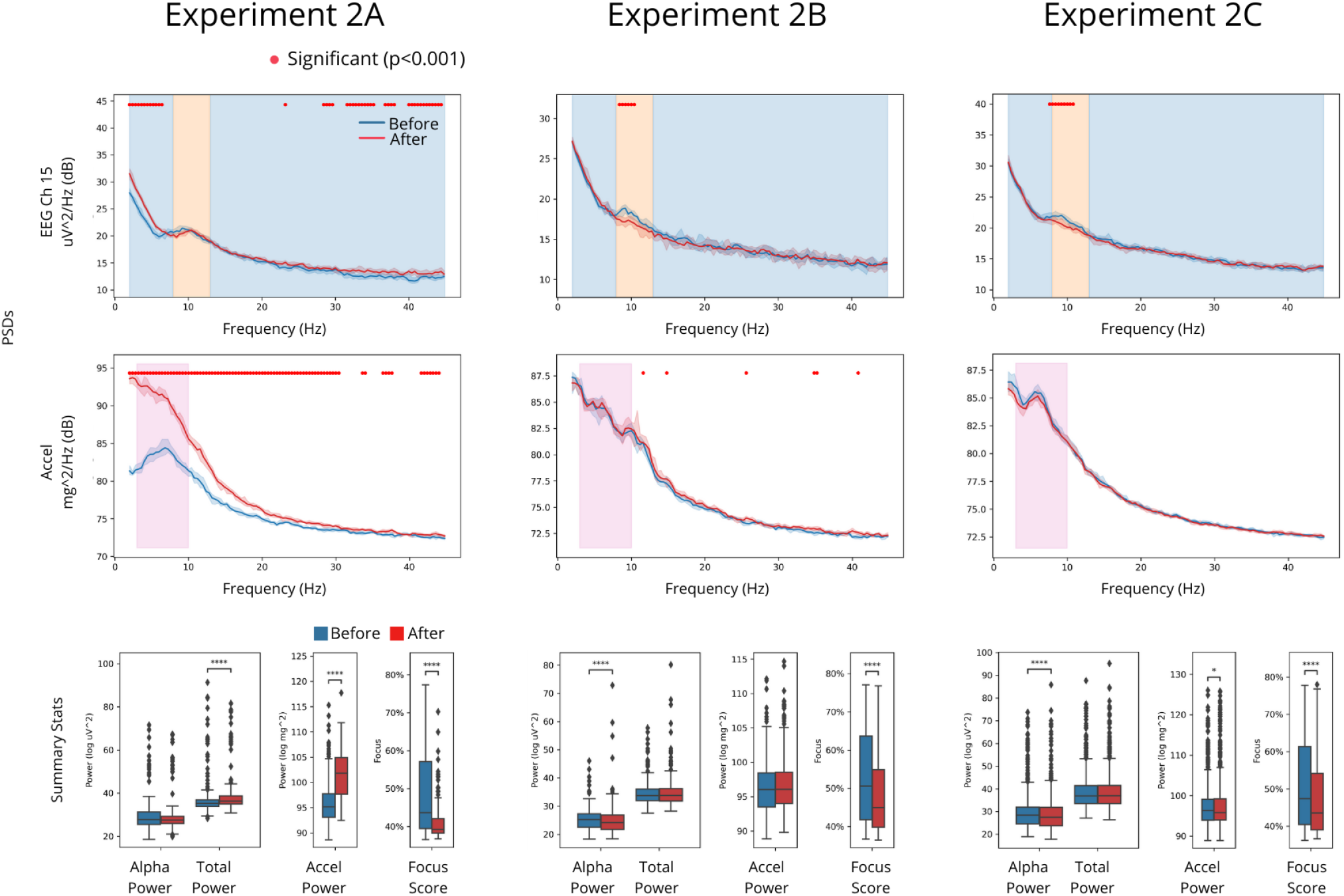
Focus score is resilient to EMG artifacts at the group level: Top row shows differences in the EEG PSDs of channel 15 after distraction relative to before. Alpha power (orange shaded region) decreases significantly in Experiments 2B and 2C, whereas Experiment 2A experiences a broadband increase in power post-distraction. Second row shows accelerometer PSDs. Experiment 2A exhibits an increase in accelerometer activity, suggesting that motion and/or EMG might underlie the broadband increase in EEG power. Experiments 2B and 2C, however, show little changes in accelerometer activity, suggesting that these experimental protocols successfully controlled for motion artifacts. Bottom row shows PSD summary statistics for alpha power (integral over the orange band in above PSDs), total power (blue band + orange band), accelerometer power (pink band, 3-10 Hz), and focus score. Before/after PSDs are calculated using data from -4s to 0s, and 0s to 4s respectively, with t=0 corresponding to the distraction event. Each data point represents the median + 95% confidence interval of N distraction events pooled together across sessions, where N=192, 207, and 505 respectively. Significance was tested using Wilcoxon signed-rank test. For line plots, alpha=0.001. For box plots, * corresponds to p<0.05, ** to p<0.01, *** to p<0.001, and **** to p<0.0001.

### 7.3 Group analysis shows EEG response is not due to auditory evoked potentials

Finally, in Figure 16, we expanded the spectral analysis in Figure 15 to show changes in activity as a function of time. This analysis was performed at the group level, with distraction onset time being t = 0 s. We found that the drop in focus and corresponding alpha suppression observed in Figure 15 is a sustained phenomenon that lasts many seconds post-distraction, rather than the brief transient one would suspect to be associated with an auditory evoked potential (e.g., response to the Slack notification). This can be seen in the top two rows of Figure 16. Additionally, we note that for Experiments 2B and 2C, there are no increases in total power or accelerometer activity associated with the distraction onset, reaffirming that these experiments do not induce any motion artifacts. Interestingly, while Experiment 2A shows an increase in alpha coincident with the spike in total power and accelerometer power (peaking at 1.8 s), channel 6 subsequently shows a significant drop in alpha power (3.5 s onwards). This suggests that Experiment 2A might induce a drop in alpha power, but this is masked by the broadband upshift associated with motion artifacts.

**Figure 16.**
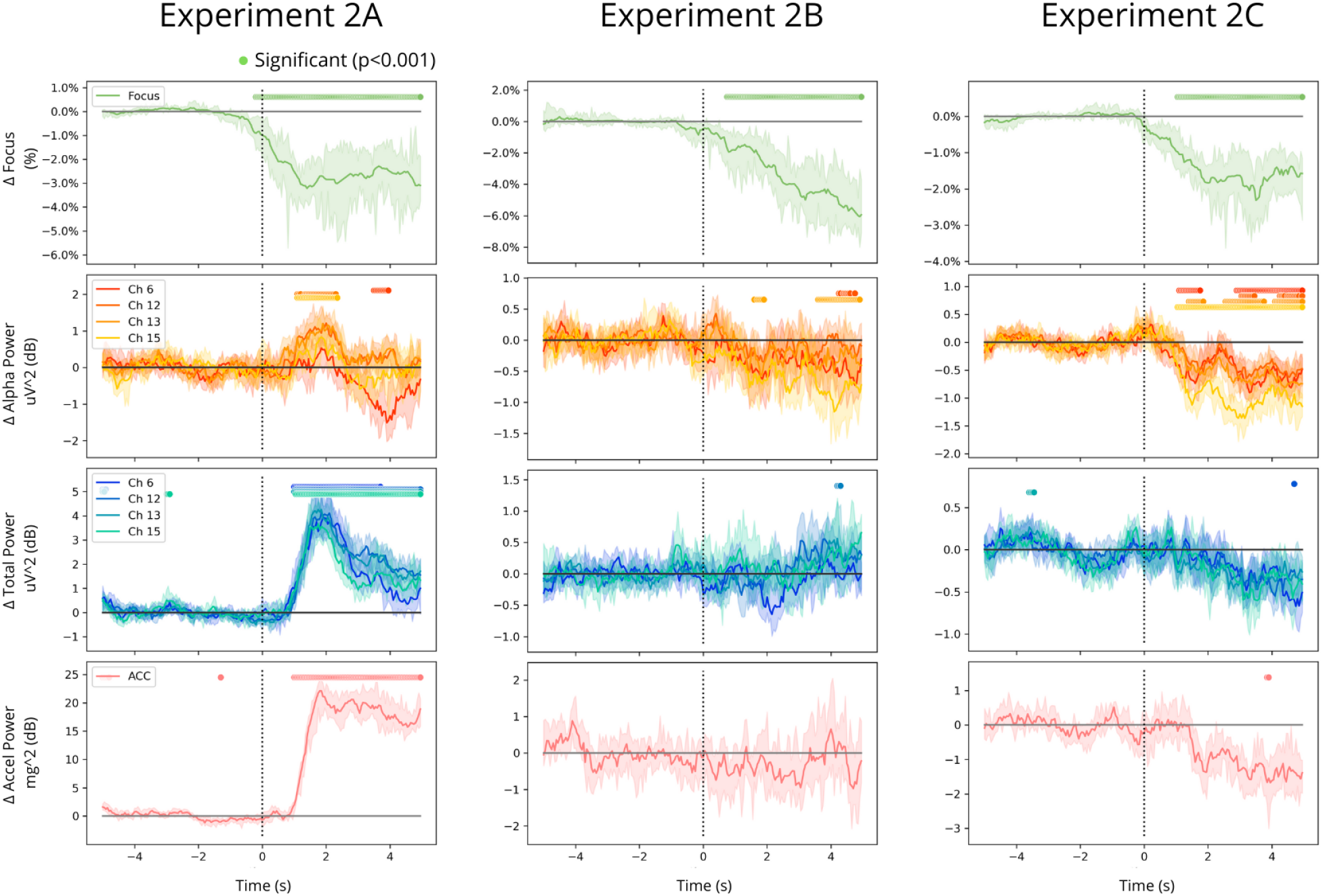
Algorithm response is not driven by auditory evoked potentials: Shown here are changes in the predicted focus level (1st row) and spectral powers (rows 2, 3, and 4) as a function of time, with t=0 s corresponding to the distraction notification. As seen in Figure 15, Experiments 2B and 2C show significant decreases in alpha power (2nd row), which drives the algorithm’s drop in predicted focus post-distraction (1st row). Given that the EEG alpha changes persist for as long as 4 seconds post-distraction, it is unlikely that they represent an auditory evoked response but, rather, reflect the user’s change in behavior. Rows 2, 3, and 4 correspond to orange, blue+orange, and pink bands in Figure 15, respectively. Data are baseline subtracted such that the data from -4s to 0s has a median of zero. Each data point represents the median + 95% confidence interval of N distraction events pooled together across sessions, where N=192, 207, and 505 respectively. Significance was tested using Wilcoxon signed-rank test with alpha=0.001.

## 8 Performance during daily use

To validate the consistency and repeatability of the model in identifying distractions, we analyzed the data of the same participant who has undergone Experiment 2A on three consecutive days (Figure 17). The results are consistent with our previous results confirmed in a cohort of participants: our focus measure consistently drops after each distraction across all days. Further, QA scores of 82.7%, 86.2% and 88.4% confirm the consistency of the model on the same person and its reliability for daily use.

**Figure 17.**
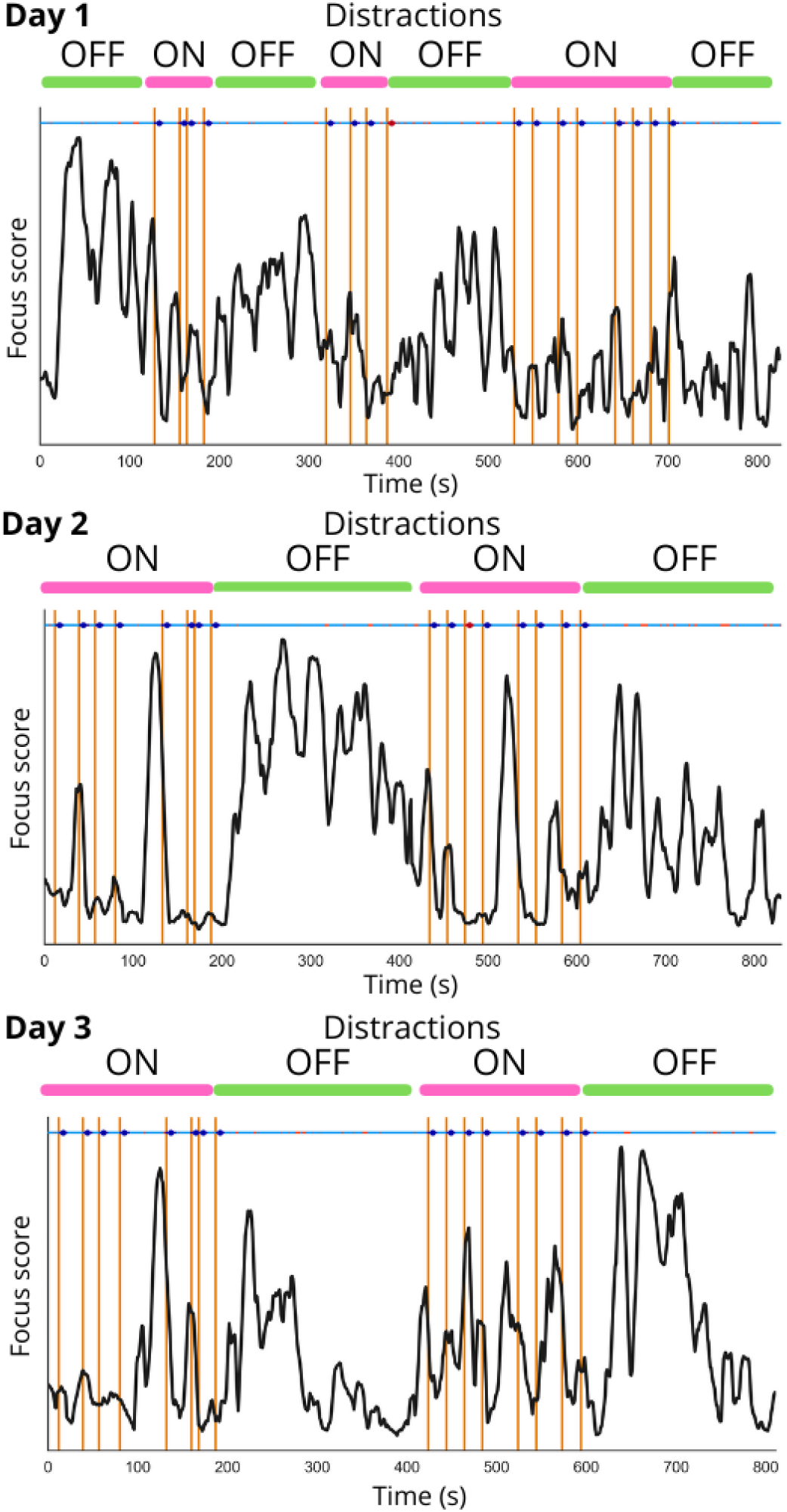
Model prediction over multiple days. Model prediction (black) and distraction timestamps (orange) of a single participant over three consecutive days undertaking Experiment 2A. The respective QA scores were 82.7% (day 1), 86.2% (day 2) and 88.4% (day 3)

To further test the generalization of the model, we examined its performance longitudinally across data of multiple people recorded over multiple sessions. These subjects completed all three experiments on different days. On average there was a gap of 10 days between each recording of three experiments’ configurations. So far, we were able to assess this on 6 subjects including the one discussed above. Table 2 summarizes the performance scores on each of these subjects.

**Table 2.** QA scores for six participants who have done all three experiments.

| Participant ID | Experiment 2A | Experiment 2B | Experiment 2C |
| --- | --- | --- | --- |
| 1 | 79.5% | 84.5% | 77.6% |
| 2 | 82.9% | 83.2% | 67.3% |
| 3 | 84.0% | 87.7% | 87.7% |
| 4 | 86.9% | 73.3% | 74.5% |
| 5 | 74.4% | 75.9% | 84.3% |
| 6 | 77.7% | 74.9% | 86.0% |
| Average | 80.9% | 79.9% | 79.6% |

The results in Table 2 show that our model provides a robust and repeatable measure of focus and is able to detect embedded distractions. The average QA scores in all the experiments are about 80%, which means our model is generalizable with a consistent prediction of distractions independent of the subject, time and conditions. This also validates the capability of the algorithm for daily use with minimal amount of training.

## 9 Discussion

In the past five years, Neurable’s effort has been directed towards building a wearable EEG headset in the form of high-quality wireless headphones (Enten) and developing a cutting-edge analytics platform to provide insights into the brain activity of a user. Enten uses dry EEG electrodes made of fabric that are integrated into the ear pads, making them invisible to external users and simplifying the setup of the EEG device. Neural data are transmitted wirelessly via Bluetooth to a proprietary analytic platform that uses state-of-the-art signal processing and machine learning algorithms to provide an estimate of the user’s focus in real-time.

Noninvasive techniques for measuring neural activity like EEG can capture much larger signals than brain activity, such as muscular movements (EMG), potentially limiting or interfering with the amount of usable neural data^17^. Traditional EEG systems use wet electrodes to increase the signal quality. When developing a new EEG device based on dry electrodes, it is critical to validate that it is capable of measuring neural activity and control for potential artifact contamination. In this research, through multiple validation steps, we have demonstrated that Enten is capable of capturing relevant brain activity of similar quality to a research-grade EEG headset. We showed that changes in behavior correspond to measurable changes in EEG captured by Neurable’s headset. Moreover, while traditional EEG headsets have full head coverage, Enten is capable of gathering insights into brain activity from sensors placed around the ears. With the development of Enten, Neurable is one step closer to the commercialization of an affordable and user-friendly EEG device for tracking focus for everyday use.

We used Enten to develop and validate a machine-learning algorithm capable of estimating the level of focus of a user in real time from EEG activity. Our algorithm uses the normalized power in the alpha band at several electrodes as features, and state-of-the-art machine learning to translate these features into a focus score. We demonstrated on a large population of 70+ users that alpha features extracted from Enten correlate with focus during a Stroop task. Moreover, we showed that distractions cause an immediate loss of focus that can be detected in real-time with EEG. With additional experiments, we further validated that our model was not driven by EMG activity or other confounds (e.g., heart rate and EOG), and that it was robust across time and users.

These promising results obtained with Enten make an important step towards building neurotechnologies for everyday use. In particular, we envision Enten to be used to enhance our productivity, filtering out distractions when we are focused and suggesting we take a break to avoid burnout. Future work will be directed towards expanding our focus algorithm to use other neural and physiological features. For example, through our analysis we showed that power in both alpha and beta EEG bands is reduced after loss of focus. Integrating beta features into our algorithm may make it more robust and generalizable. We will also use Enten to estimate the focus score in other daily tasks, to verify the generalizability of our findings to other domains.

In this white paper, we considered the difficulty of the task at hand being a proxy to measure the level of focus of the user. The rationale behind this choice was that when the task is difficult – either for incongruence between color and word (as in Stroop) or the presence of an auditory distraction that requires us to attend another task – our brain does not have enough resources to dedicate to the main task. Suppression of alpha activity when tackling difficult tasks – as we found in our experiments – has been previously reported in a variety of other tasks^18^. To make our findings easily accessible to any reader, we used the word “focus” as the outcome estimated by our ML algorithm. Nevertheless, our findings could also be interpreted in the context of cognitive load and mental effort. In fact, lower alpha activity in the temporal cortex has been associated with increased cognitive load^18,19^.

## 10 Conclusion

In summary, Enten represents a leap towards bringing neurotechnology out of research labs to daily life, and solves several important challenges that BCIs face. First, Enten uses dry electrodes made of fabric to record EEG activity, hence simplifying setup and comfort for the user. Second, EEG sensors are integrated in a device of everyday use, a pair of high-quality headphones, making them unnoticeable to an external user. Third, EEG data from several sensors are transmitted wirelessly with millisecond accuracy and processed in real-time to extract neural features. Fourth, machine learning models accurately estimate focus from neural activity recorded around the ears (temporal lobe). Finally, Enten provides robust and statistically-validated focus estimation across users, days, and experimental settings. With Enten, Neurable has built a multiuse, everyday BCI that could be used to gather insight into our brain, as well as a novel interface to control other devices with our brain activity.

## Acknowledgment

We would like to thank all participants in our experiments, the incredible Neurable team, and all our advisors and investors for having made this research possible.

## Supplementary Figures

Included here are supplementary figures for the papers, which provide additional characterization of Enten eyes-closed alpha response and signal quality.

**Figure 18.**
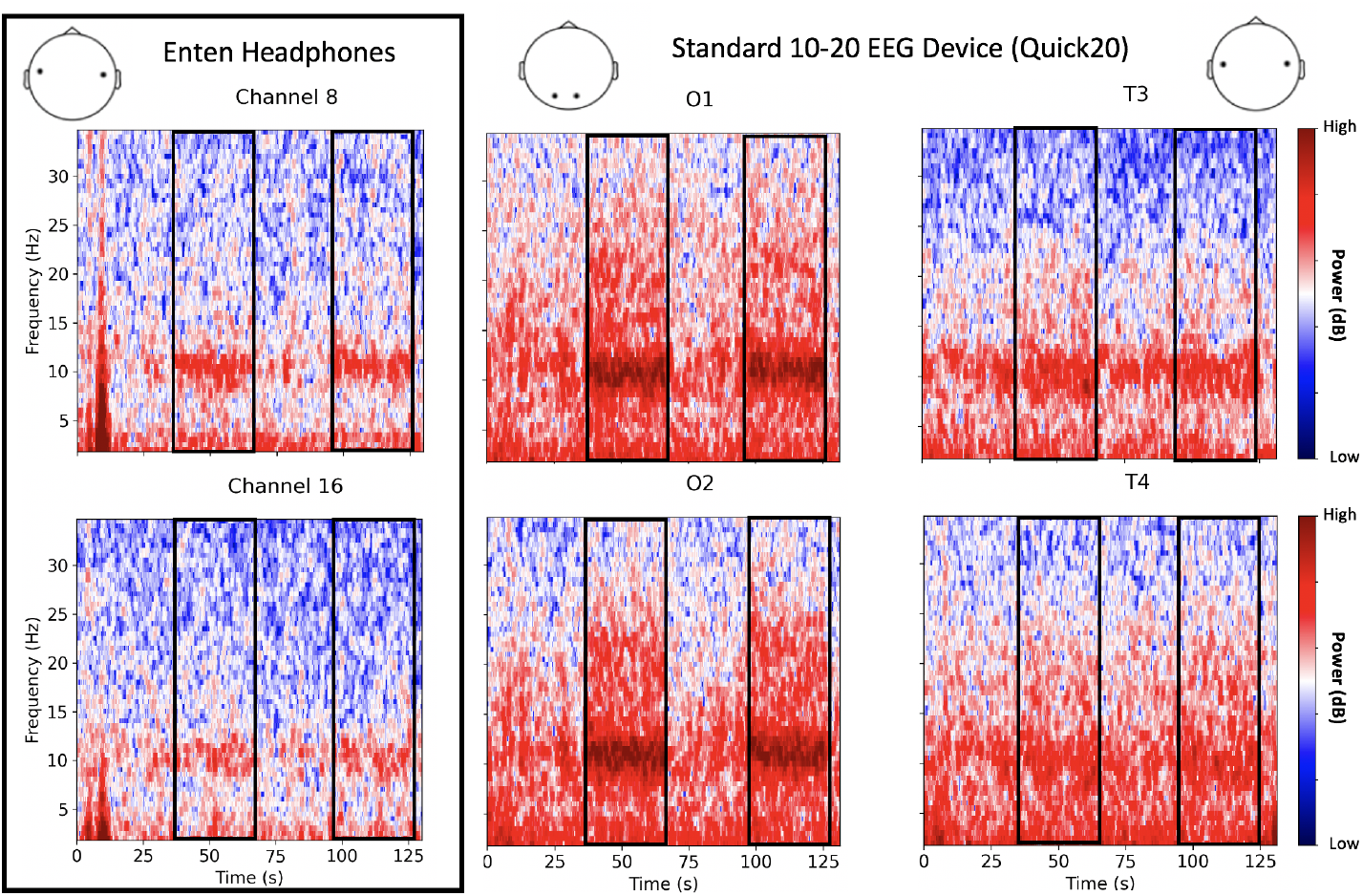
Spectrogram of eye opened and closed of a different participant, as recorded using Enten (left) and Quick20 (middle and right). 2 minutes experiment: eye opened for 30 seconds followed by 30 seconds eye closed, and repeated the procedure twice. The mean difference of the PSD between eye closed and opened are 3.08 dB, 2.94 dB, 9.71 dB, and 6.22 dB for T3, T4, O1, and O2 respectively (left and middle). The mean difference of the PSD are 4.12 dB and 5.65 dB for Channel 8 and Channel 16 respectively (right).

**Figure 19.**
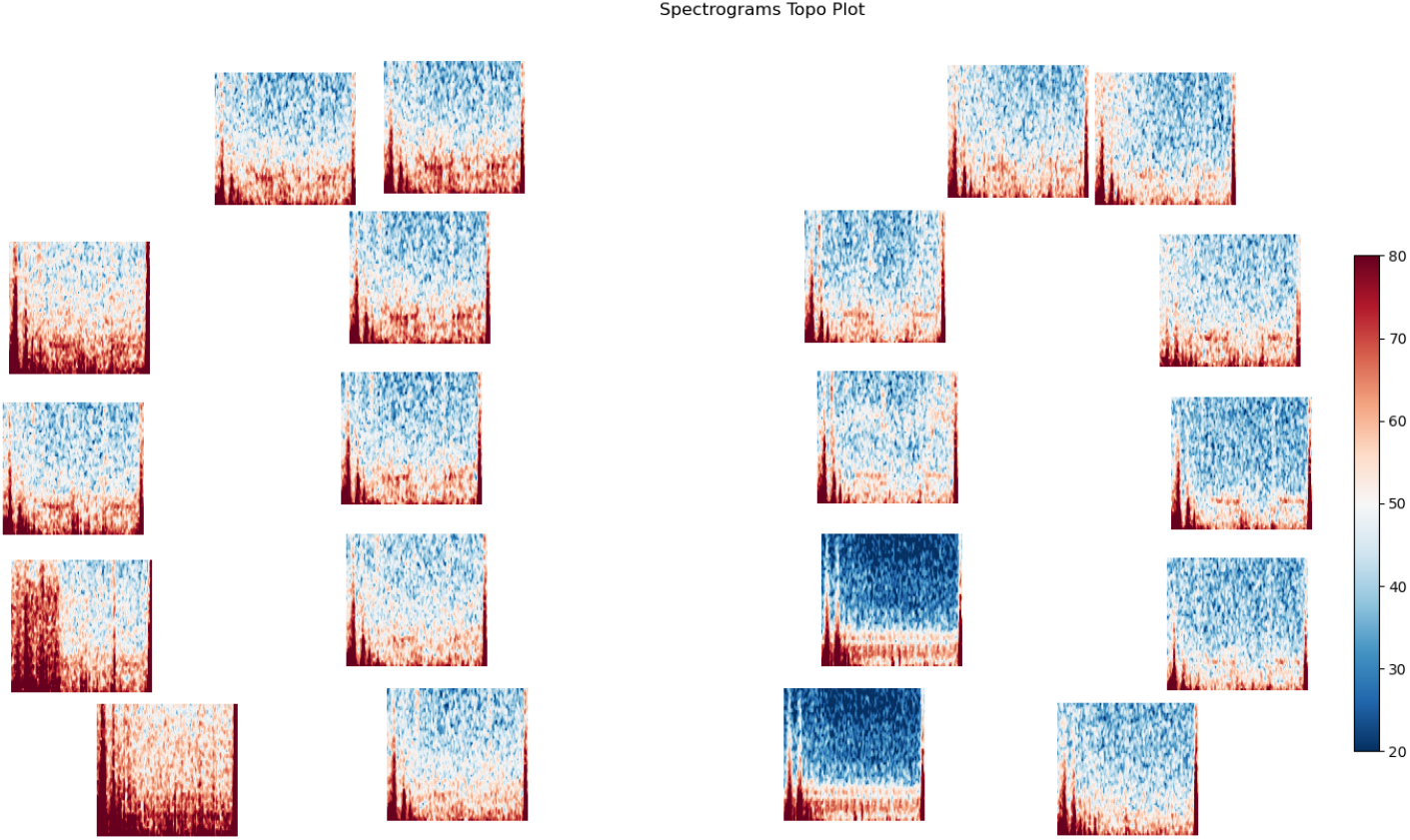
Alpha response for each electrode pf the Enten headphones. Participants had their eyes open and closed for 30 seconds each. This procedure was repeated twice

**Figure 20.**
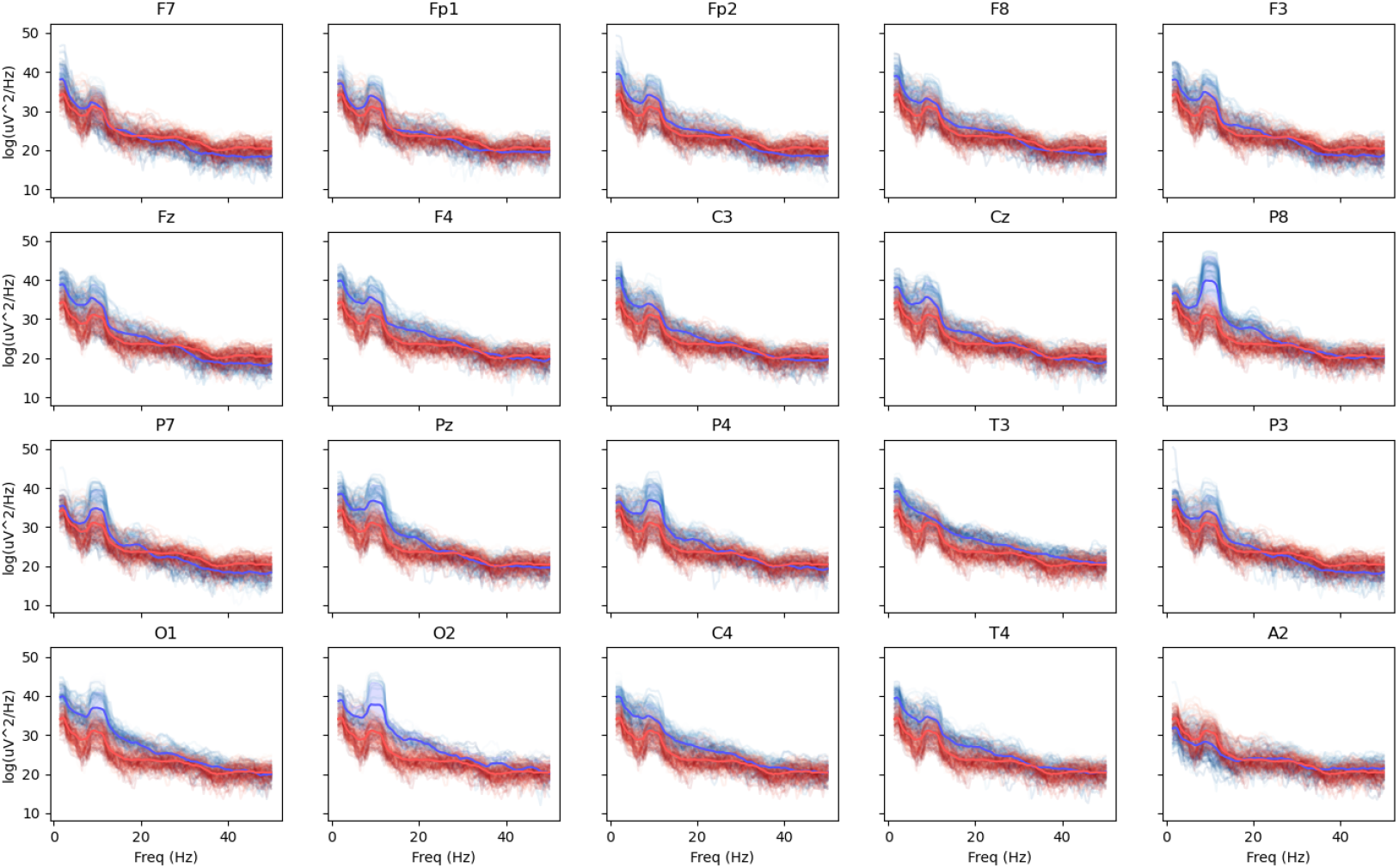
Headset comparison by channel. Eyes closed alpha for Enten Channel 17 (red) versus a commercially available 10-20 system (Cognionics Quick20, blue). Same subject in both cases. PSDs are calculated based on 100 seconds of eyes closed data. Translucent lines indicate individual epochs (2 second duration, 50% overlapping), and solid lines denote the average across these epochs.

**Figure 21.**
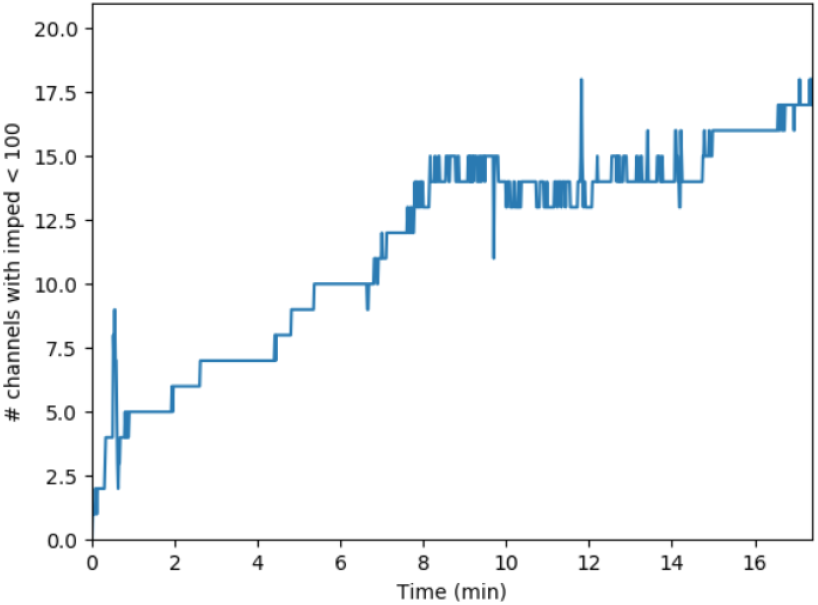
Number of channels with low impedance (<100 kΩ) at as a function of time. Electrodes require several minutes to settle. This informed us to require subjects to wear the headphones for a few minutes before starting the experiment.

## Notes

### Competing Interest Statement

The authors have declared no competing interest.

